# Overcoming the Reproducibility Crisis - Results of the first Community Survey of the German National Research Data Infrastructure for Neuroscience

**DOI:** 10.1101/2022.04.07.487439

**Authors:** Carsten M. Klingner, Michael Denker, Sonja Grün, Michael Hanke, Steffen Oeltze-Jafra, Frank W. Ohl, Janina Radny, Stefan Rotter, Hansjörg Scherberger, Alexandra Stein, Thomas Wachtler, Otto W. Witte, Petra Ritter

**Author notes:** Corresponding authors: Carsten Klingner;, Petra Ritter.

## Abstract

The lack of reproducibility of research results is a serious problem – known as “the reproducibility crisis”. The German National Research Data Infrastructure (NFDI) initiative implemented by the German Research Foundation (DFG) aims to help overcoming this crisis by developing sustainable solutions for research data management (RDM). NFDI comprises domain specific consortia across all science disciplines. In the field of neuroscience, NFDI Neuroscience (NFDI-Neuro) contributes to the strengthening of systematic and standardized RDM in its research communities. NFDI-Neuro conducted a comprehensive survey amongst the neuroscience community to determine the current needs, challenges, and opinions with respect to RDM. The outcomes of this survey are presented here. The German neuroscience community perceives barriers with respect to RDM and data sharing mainly linked to (1) lack of data and metadata standards, (2) lack of community adopted provenance tracking methods, 3) lack of a privacy preserving research infrastructure for sensitive data (4) lack of RDM literacy and (5) lack of required time and resources for proper RDM. NFDI-Neuro aims to systematically address these barriers by leading and contributing to the development of standards, tools, and infrastructure and by providing training, education, and support, as well as additional resources for RDM to its research community. The RDM work of NFDI-Neuro is conducted in close collaboration with its partner EBRAINS AISBL, the coordinating entity of the EU Flagship Human Brain Project, and its Research Infrastructure (RI) EBRAINS with more than 5000 registered users and developers from more than 70 countries of all continents. While NFDI-Neuro aims to address the German national needs, it closely aligns with the international community and the topics of the Digital Europe Program and EU Data Spaces.

**Significance Statement:** A comprehensive survey amongst the neuroscience community in Germany determined the current needs, challenges, and opinions with respect to standardized research data management (RDM) to overcome the reproducibility crisis. Significant deficits were pointed out concerning the perceived lack of standards for data and metadata, lack of provenance tracking and versioning of data, lack of protected digital research infrastructure for sensitive data and the lack of education and resources for proper RDM. Yet, at the same time, an overwhelming majority of community members indicated that they would be willing to share their data with other researchers and are interested to increase their RDM skills. Thus, the survey results suggest that training, the provision of standards, tools, infrastructure and additional resources for RDM holds the potential to significantly facilitate reproducible research in neuroscience.

## Introduction

It is well acknowledged by the research community that poor reproducibility of research results is a serious challenge – known as “the reproducibility crisis” – that hinders growths of knowledge and innovation on the one hand and leads to inefficient use of resources on the other hand (Baker, 2016; Crook et al., 2020; Loss et al., 2021; Poldrack et al., 2019; Stodden et al., 2016). The German National Research Data Infrastructure Initiative (NFDI) implemented by the German Research Foundation (DFG) has allocated €900 million over the course of 10 years to foster research data management (RDM) across all research domains in Germany with the aim of overcoming the reproducibility crisis. NFDI comprises domain specific consortia across all science disciplines. In the field of neuroscience, the consortium NFDI Neuroscience (NFDI-Neuro; https://nfdi-neuro.de) has started to closely interact with their community to overcome the challenges in RDM (Ebert et al., 2021; Denker et al., 2021a, 2021b; Hanke et al., 2021; Klingner et al., 2021; Wachtler et al., 2021). The NFDI-Neuro initiative is closely aligned to core topics of the Digital Europe Program, including artificial intelligence, cybersecurity, supercomputing, and the European Health Data Space. It aims to offer next generation RDM solutions and corresponding training and education opportunities for the whole research community. NFDI-Neuro consists of 10 applying German research institutions, 15 participating institutions and 20 participating scientists – geographically distributed across whole Germany (Figure 1).

**Figure 1:**
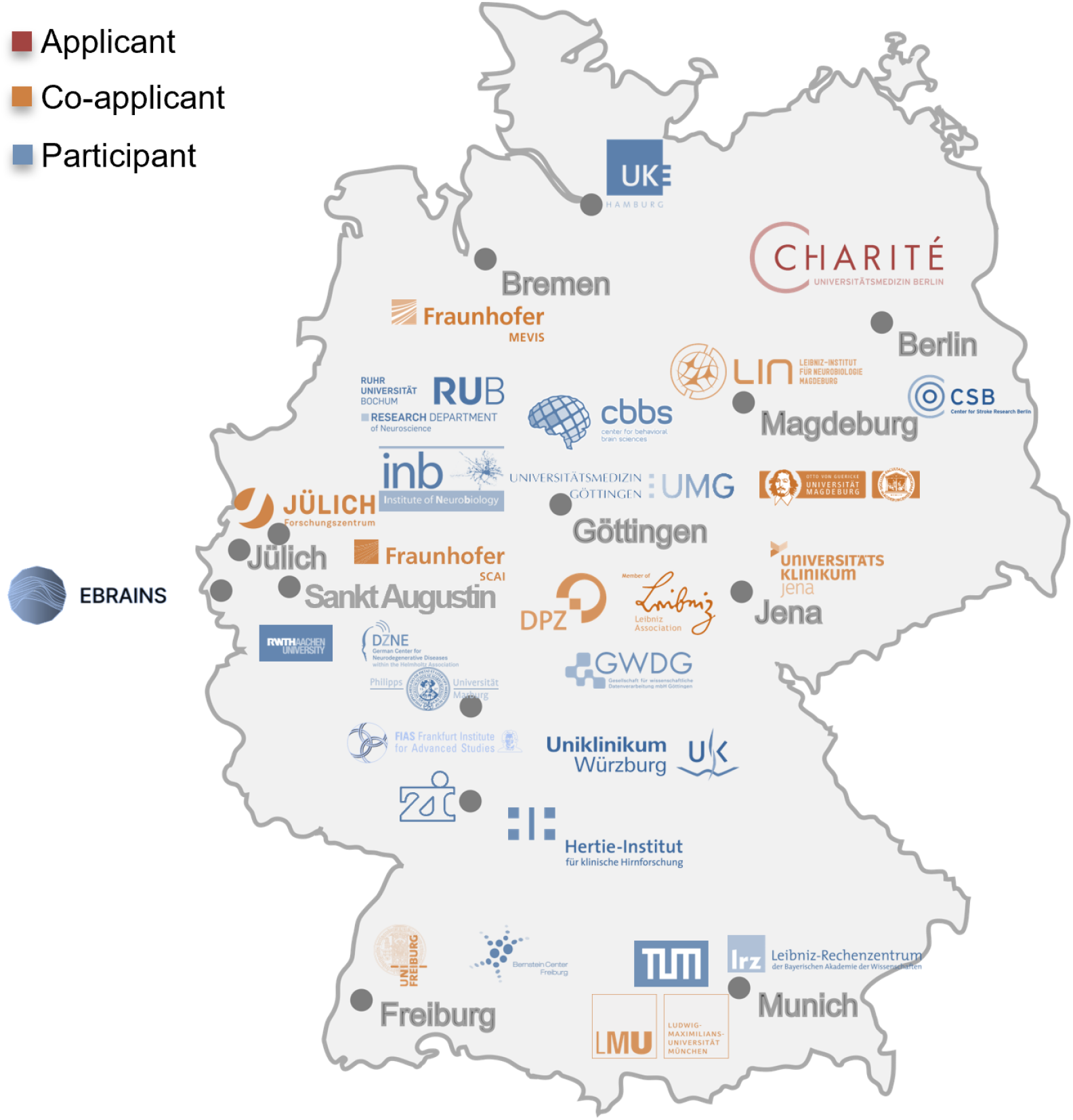
NFDI-Neuro covers a geographically distributed diverse Neuroscience community and collaborates with the international partner EBRAINS AISBL.

Amongst the participating entities are the German Neuroscience Society with 2300 members, the Bernstein Network Computational Neuroscience with 400 members, the German Society for Clinical Neurophysiology and Functional Imaging with 4000 members, and EBRAINS AISBL – the coordinator of the EBRAINS (ebrains.eu) Research Infrastructure (RI) with more than 5000 registered users from 75 countries and 900 institutions. EBRAINS AISBL is also the coordinating entity of the EU Flagship Human Brain Project (Amunts et al., 2016, 2019, 2022; Amunts, Katrin et al., 2022; Bjerke et al., 2018; Quaglio et al., 2017; Salles et al., 2019; Schirner et al., 2022; Tiesinga et al., 2015) with currently 134 partner institutions from more than 20 countries. The EBRAINS RI under lead of EBRAINS AISBL has recently been accepted to the European Strategy Forum for Research Infrastructures (ESFRI) roadmap securing its long-term persistence over the next decades as a reference RI for the international neuroscience community. This and other international brain initiatives aim to use computational models as theoretical frameworks and atlases as spatial anchors for multi-scale neuroscience data integration – to turn data into knowledge and understanding (Amunts et al., 2013, 2020; Bjerke et al., 2018; Ritter et al., 2013; Schirner et al., 2018). EBRAINS RI also provides digital workflows and provenance tracking (e.g. (Schirner et al., 2015; Wagner et al., 2021) to enable research reproducibility.

Historically, the neuroscience community has been spearheading developments for digital RDM and computational workflows as reflected by the foundation of the International Neuroinformatic Coordination Facility (INCF, cf., (Abrams et al., 2021; Bjaalie and Grillner, 2007; Poline et al., 2022) in 2005 and the EU flagship Human Brain Project starting in 2013 with its e-infrastructure EBRAINS. These initiatives share the common goal of developing a unified and systematic understanding of brain function in health and disease by integrating data and knowledge from the various subdisciplines. NFDI-Neuro will tackle the conceptual and logistic challenge of the integration and standardized representation of data and metadata and practically make these emerging solutions and infrastructures available to neuroscientists for use in their daily work by offering training and support. In this way, NFDI-Neuro aims to foster the reproducibility of research and leveraging computational neuroscience as data integrating discipline that transforms data into knowledge and understanding.^1234^

To obtain a good understanding of the present RDM situation in the neuroscience community, NFDI-Neuro conducted a community survey with the focus on the following questions: What are the largest obstacles and most pressing needs perceived by the neuroscience community? How does the community self-assess its present RDM proficiency? The survey was preceded by five community workshops conducted between 2019 and 2021 and organized by NFDI-Neuro, an ongoing monthly webinar series (https://webinar.nfdi-neuro.de) dedicated to the topic of RDM in neuroscience to increase awareness and engagement among members of the German neuroscience community and a weekly or biweekly community meeting enabling exchange about RDM topics and updates on latest developments^5^. Here, we report and analyse the results of the survey – the outcome of which was also used for shaping the specific workplan of the NFDI-Neuro consortium.

## Methods

The survey was developed based on an existing RDM survey by the partner consortium NFDI4Bioimage^6^. It was adapted by the NFDI-Neuro team to address questions specific to the neuroscience research domain. The present survey comprises 20 sets of questions, where some sets contain multiple questions resulting in a total of 114 questions presented to each survey participant. The survey takes on average 10-15 minutes when all questions are answered. We used the tool LimeSurvey (https://www.limesurvey.org/de/) and conducted the survey in compliance with the EU General Data Protection Regulations (GDPR) as approved by the institutional data protection officer (DPO). The online questionnaire was made available for two months between Sept-1^st^, 2021 and Nov-1^st^, 2021 via the website of the NFDI-Neuro initiative (https://nfdi-neuro.de/). We announced the survey via several channels, including email lists of the German neuroscience communities, such as the German Society for Clinical Neurophysiology, the German Neuroscience Society (GNS), and the Bernstein Network Computational Neuroscience, as well as the consortium’s mailing list and Twitter channel (https://twitter.com/NFDI_Neuro). In total 218 individuals participated in the online survey. Of those, 85 participants did not answer all questions. We included in our analysis all given answers – including those of the not fully completed questionnaires. For the data analysis and the generation of the figures we used the software package R (version 4.1.2 [“Bird Hippie”]). The survey and related collected data, as well as all analysis scripts are available publicly^7^ and can be used under public license. ^7^ https://gin.g-node.org/NFDI-Neuro/SurveyData (https://doi.org/10.12751/g-node.w5h68v)

## Results

In the following we provide all survey questions and corresponding response statistics.

### Participants represent a broad range of neuroscience disciplines

The professional position of the survey participants shows a tendency towards higher positions in the scientific hierarchy with 73 (33%) “Independent scientist and group leader / professor”, 46 (21%) “Scientists”, 56 (26%) “Student or early career researcher”, 14 (6%) “Research data management focused staff”, 6 (3%) “Tenured research staff”, 9 (4%) “Scientific support staff”, 14 (6%) “Other” (Figure 2, 22). The participants cover a wide range of neuroscience subdisciplines (selection of multiple choices possible) led by brain imaging (106 - 49%) followed by cognitive neuroscience (92 - 42%), systems and behavioral neuroscience (84 - 39%), clinical neuroscience (67 - 31%), computational/theoretical neuroscience (53 - 24%), data science (48 - 22%), neuroinformatics (31 - 14%) and cellular/molecular neuroscience (25 - 11%).

**Figure 2:**
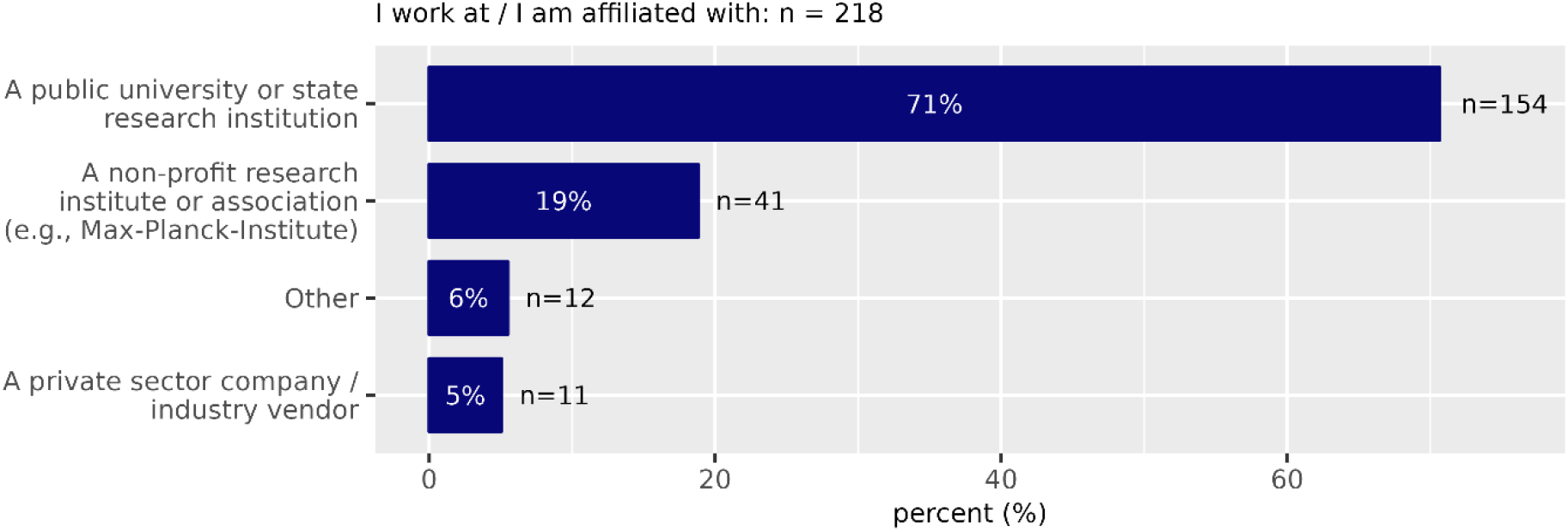
**Question 1 -** I work at / I am affiliated with Please choose **only one** of the following:

- A public university or state research institution
- A non-profit research institute or association (e.g., Max-Planck-Institute, Fraunhofer Institute, Helmholtz Center)
- Other
- A private sector company / industry vendor

**Figure 3:**
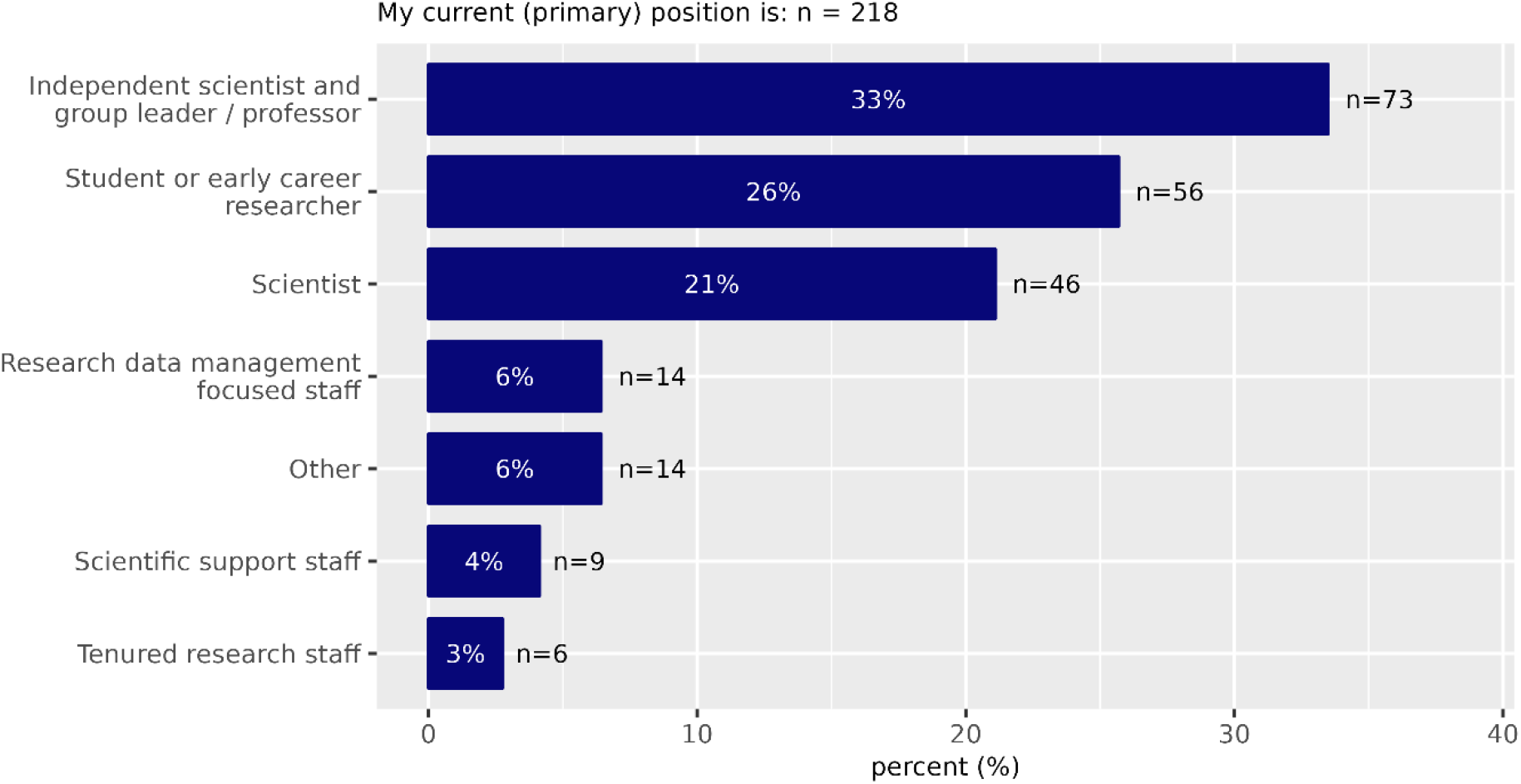
**Question 2 -** My current (primary) position is: Please choose **only one** of the following:

- Independent scientist and group leader / professor
- Student or early career researcher
- Scientist
- Research data management focused staff
- Other
- Scientific support staff
- Tenured research staff

**Figure 4:**
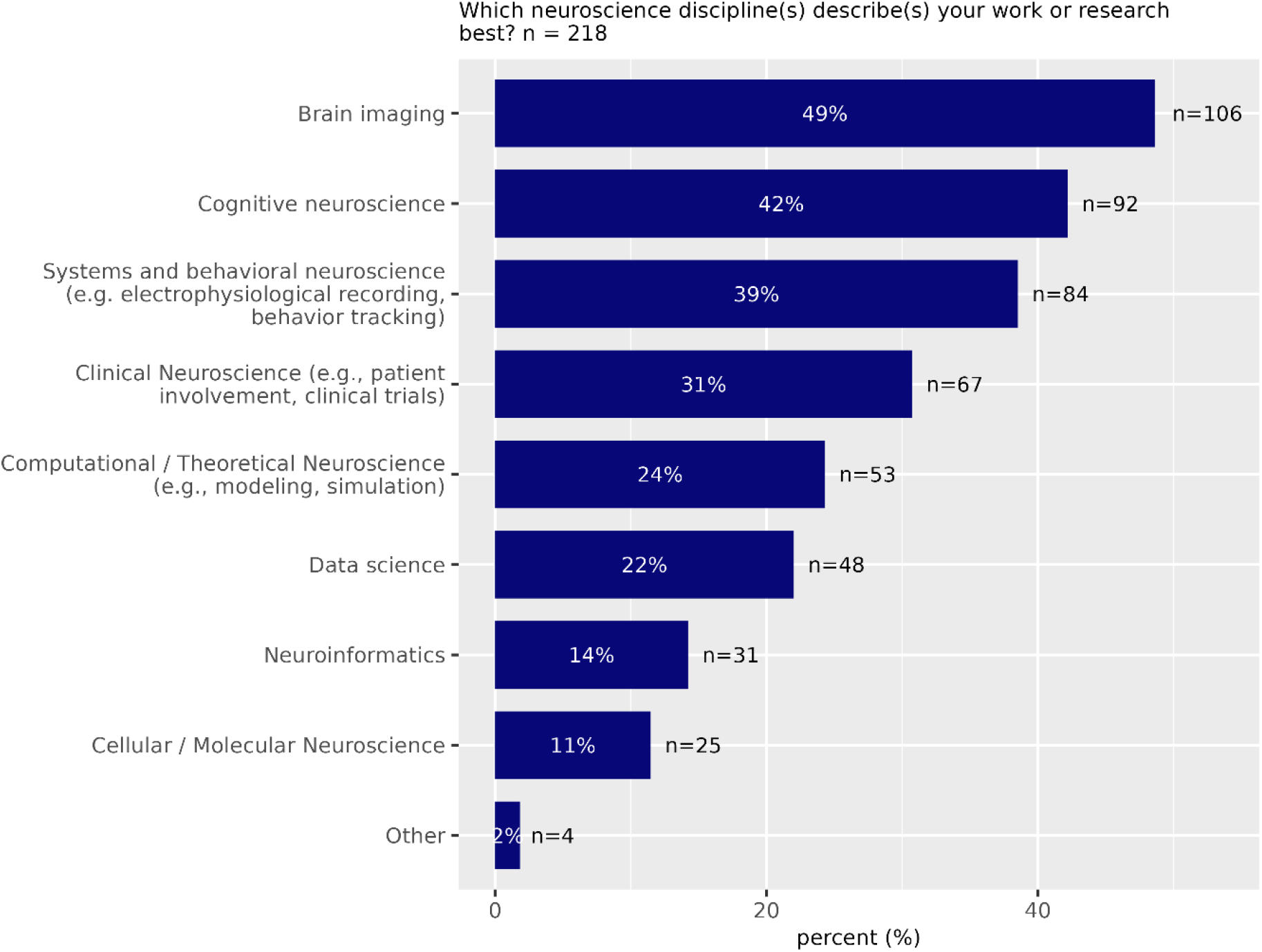
**Question 3 -** Which neuroscience discipline(s) describe(s) your work or research best? Please choose **all** that apply:

- Brain imaging
- Cognitive neuroscience
- Systems and behavioral neuroscience (e.g. electrophysiological recording, behavior tracking)
- Clinical Neuroscience (e.g., patient involvement, clinical trials)
- Computational / Theoretical Neuroscience (e.g., modeling, simulation)
- Data science
- Neuroinformatics
- Cellular / Molecular Neuroscience
- Other:

**Figure 5:**
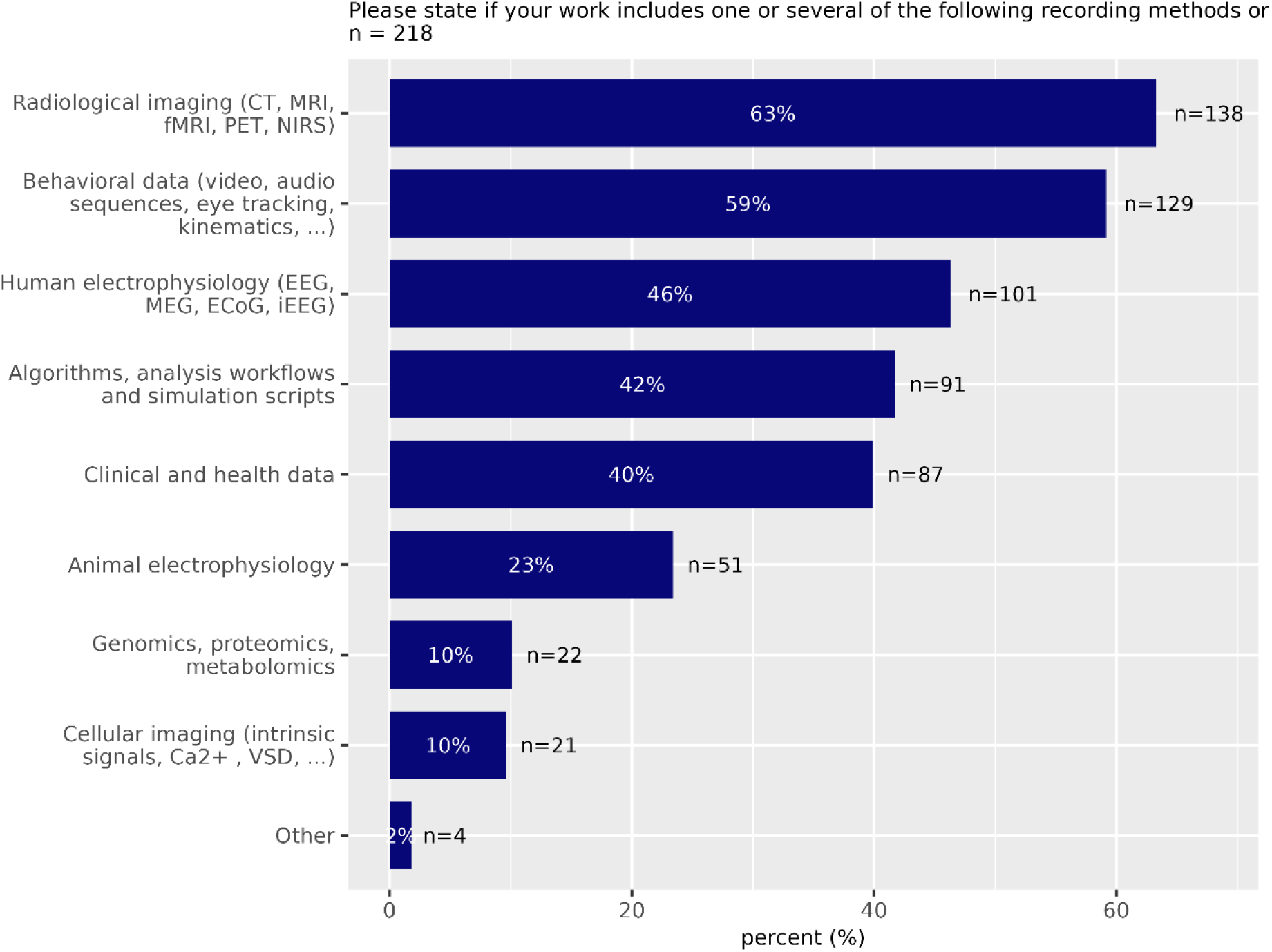
**Question 4 -** Please state if your work includes one or several of the following recording methods or data types: Please choose **all** that apply:

- Radiological imaging (CT, MRI, fMRI, PET, NIRS)
- Behavioral data (video, audio sequences, eye tracking, kinematics, …)
- Human electrophysiology (EEG, MEG, ECoG, iEEG)
- Algorithms, analysis workflows and simulation scripts
- Clinical and health data
- Animal electrophysiology
- Genomics, proteomics, metabolomics
- Cellular imaging (intrinsic signals, Ca2+, VSD, …)
- Other:

**Figure 6:**
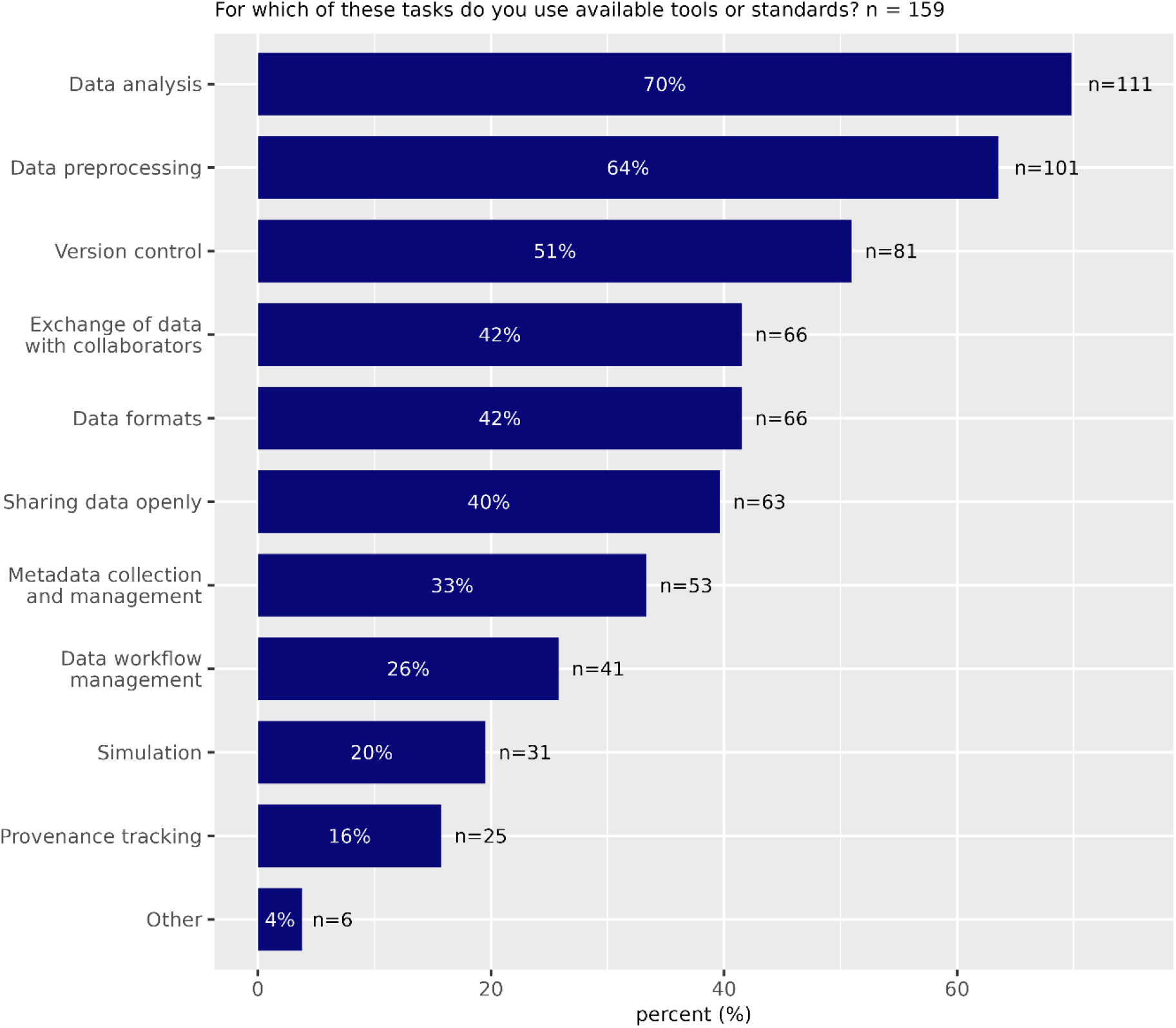
**Question 5** - For which of these tasks do you use available tools or standards? Please indicate which tools: Please choose **all** that apply and provide a comment:

- Data analysis
- Data preprocessing
- Version control
- Exchange of data with collaborators
- Data formats
- Sharing data openly
- Metadata collection and management
- Data workflow management
- Simulation
- Provenance tracking
- Other

**Figure 7:**
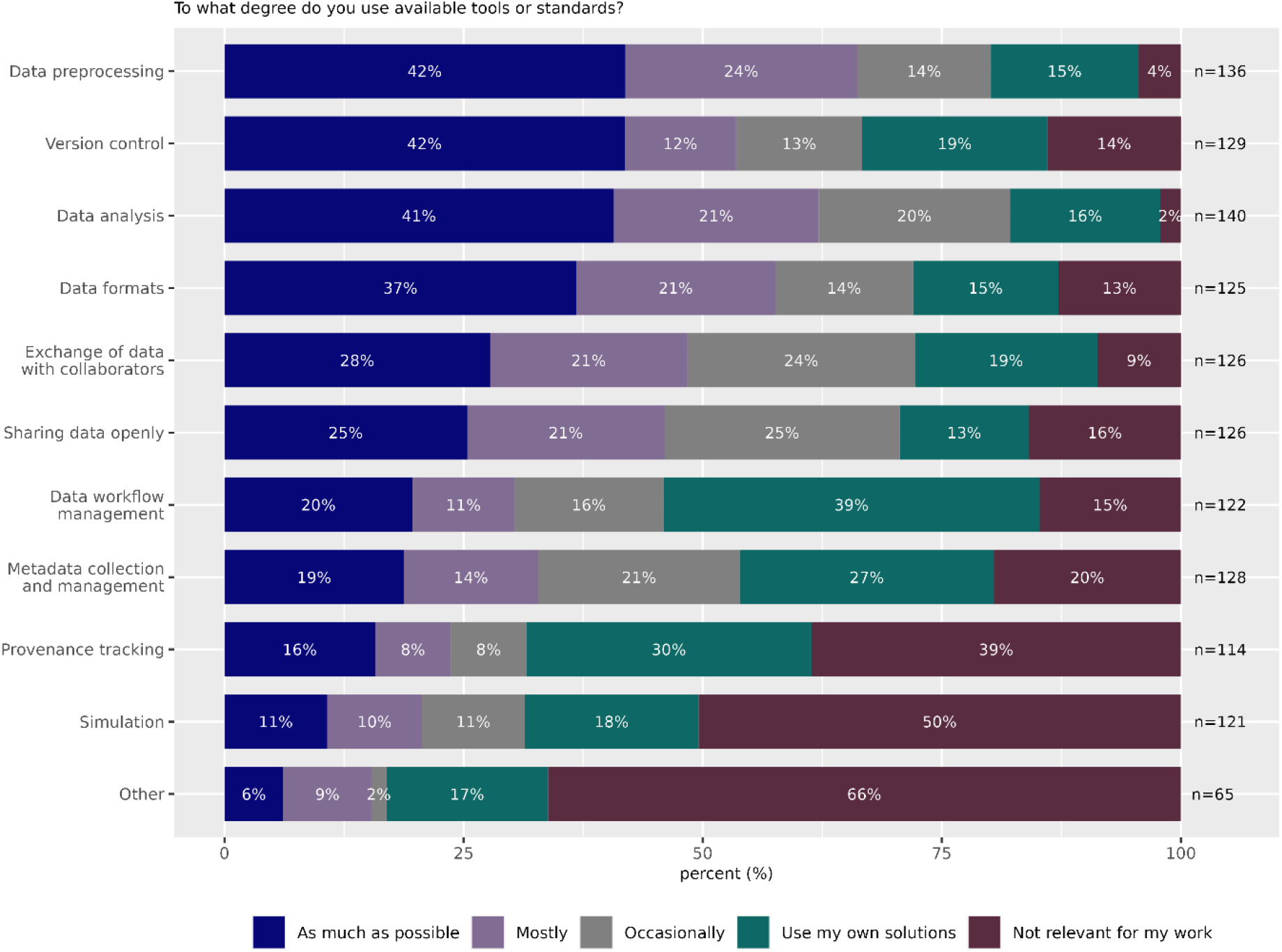
**Question 6** - To what degree do you use available tools or standards? Not at all - I use my own custom solutions / Occasionally / Mostly / As much as possible / This is not relevant for my scientific work Categories:

- Data preprocessing
- Version control
- Data analysis
- Data formats
- Exchange of data with collaborators
- Sharing data openly
- Data workflow management
- Metadata collection and management
- Provenance tracking
- Simulation
- Other

### A significant amount of research data is not yet being shared

114 (79%) of all participants share data within their institution. 95 (66%) share their data with external collaborators while only 65 (45%) share data publicly (at least one dataset). Only 13 participants (9%) had never shared any data yet (Fig. 8). A primary objective of the NFDI initiative is to improve the secondary use of data. In this context, we explored the potential availability of neuroscience data that is not yet shared publicly but is considered of general interest. We asked whether the participants own data of potential interest to other scientists for their re-use. 84 (67%) of the participants have valuable datasets available that would be useful for further exploitation, while only 20 (22%) of those participants make these data available for re-use. 76 (84%) of all participants with at least one dataset believe that other researchers could answer their research questions by re-using data from their research. However, even for this subgroup of scientists that think their data are valuable to others, 48% have never publicly shared any of their data.

**Figure 8:**
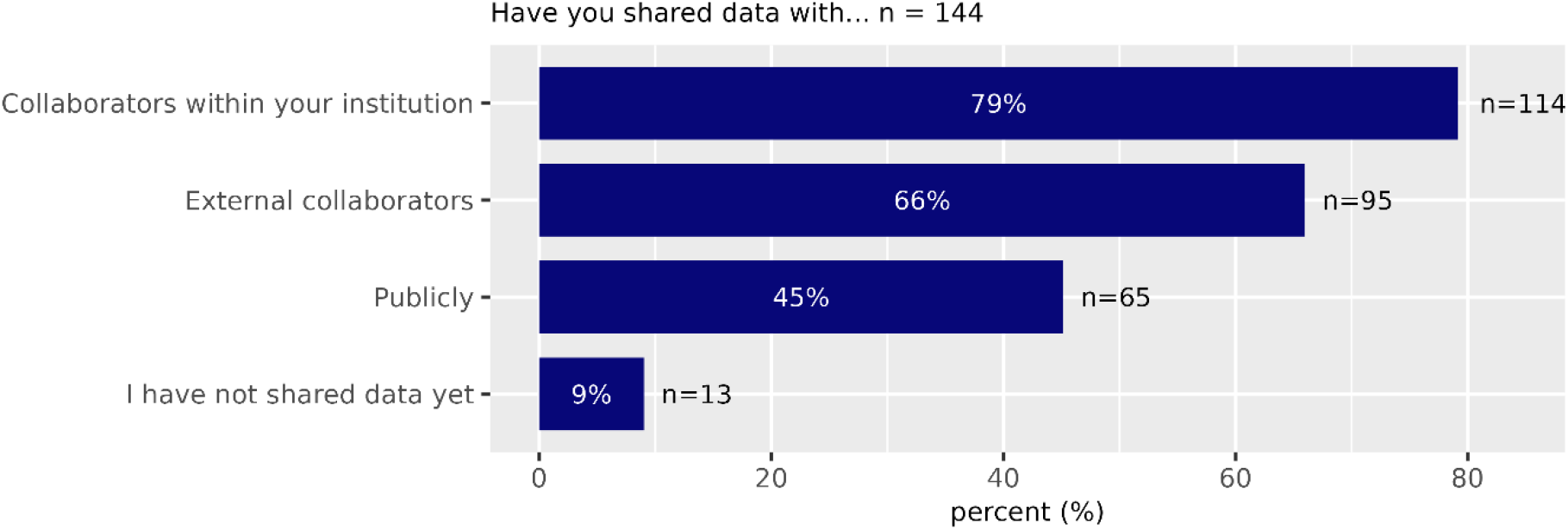
**Question 7** - Have you shared data with… Please choose **all** that apply:

- Collaborators within your institution
- External collaborators
- Publicly
- I have not shared data yet

**Figure 9:**
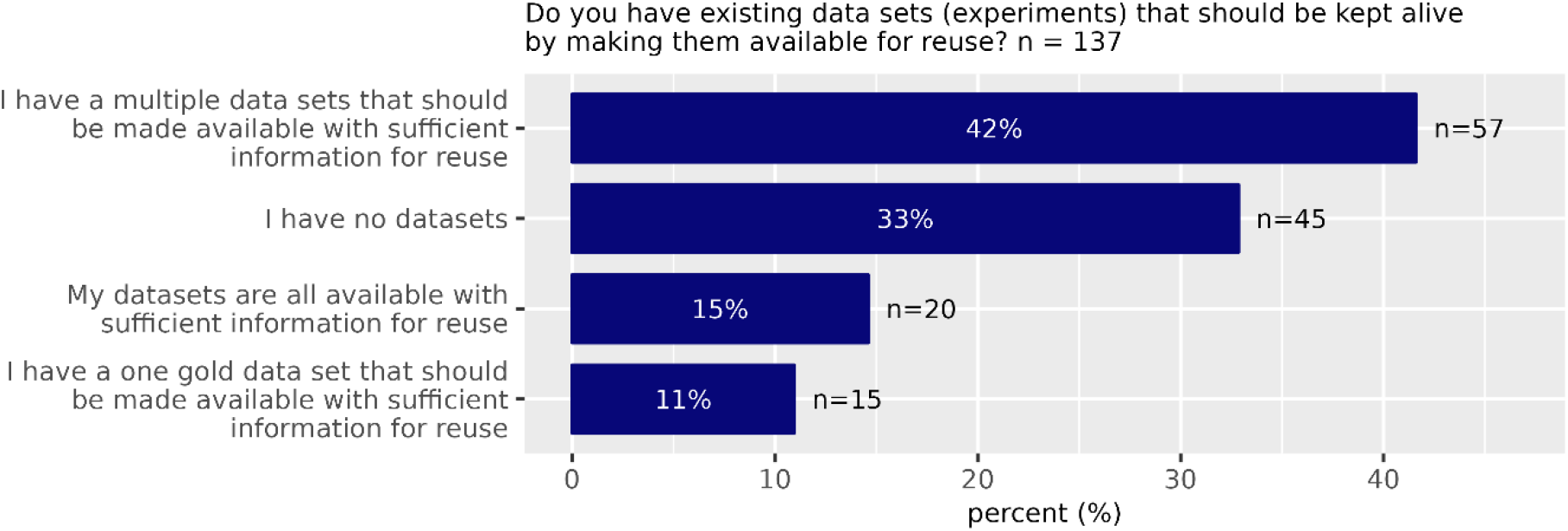
**Question 8** - Do you have existing data sets (experiments) that should be kept alive by making them available for reuse? Please choose **only one** of the following:

- I have a multiple data sets that should be made available with sufficient information for reuse
- I have no datasets
- My datasets are all available with sufficient information for reuse
- I have a one gold data set that should be made available with sufficient information for reuse

**Figure 10:**
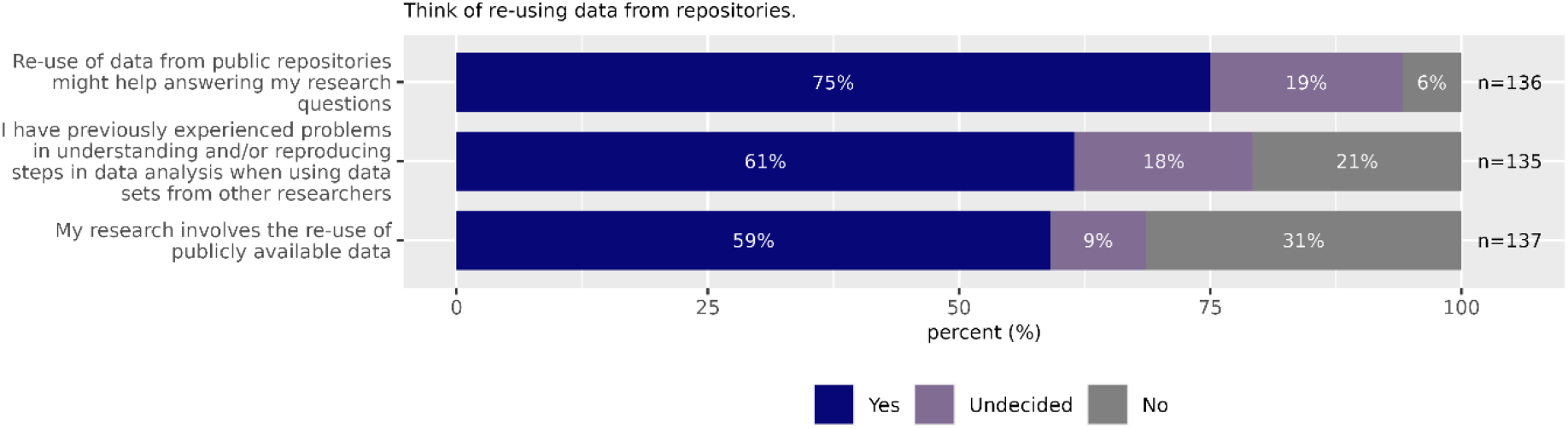
**Question 9** - Think of re-using data from repositories. Please choose the appropriate response for each item: Yes / No / Undecided

**Figure 11:**
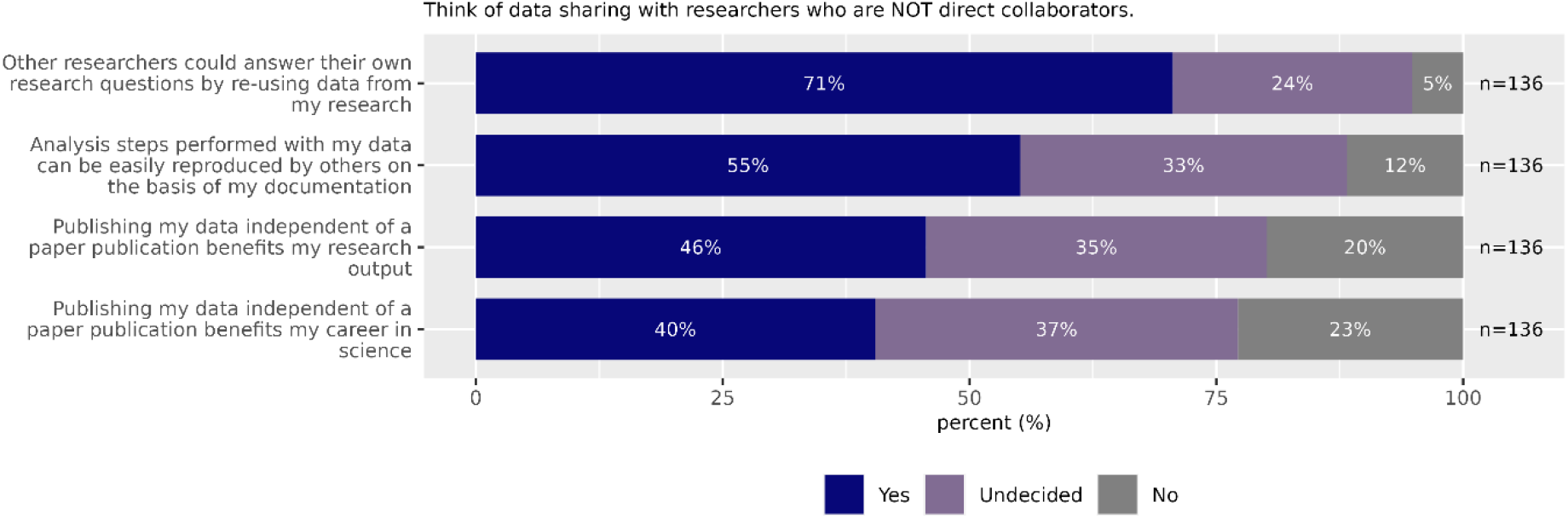
**Question 10** - Think of data sharing with researchers who are NOT direct collaborators. Please choose the appropriate response for each item: Yes / No / Undecided

- Other researchers could answer their own research questions by re -using data from my research
- Analysis steps performed with my data can be easily reproduced by others on the basis of my documentation
- Publishing my data independent of a paper publication benefits my research output
- Publishing my data independent of a paper publication benefits my career in science

**Figure 12:**
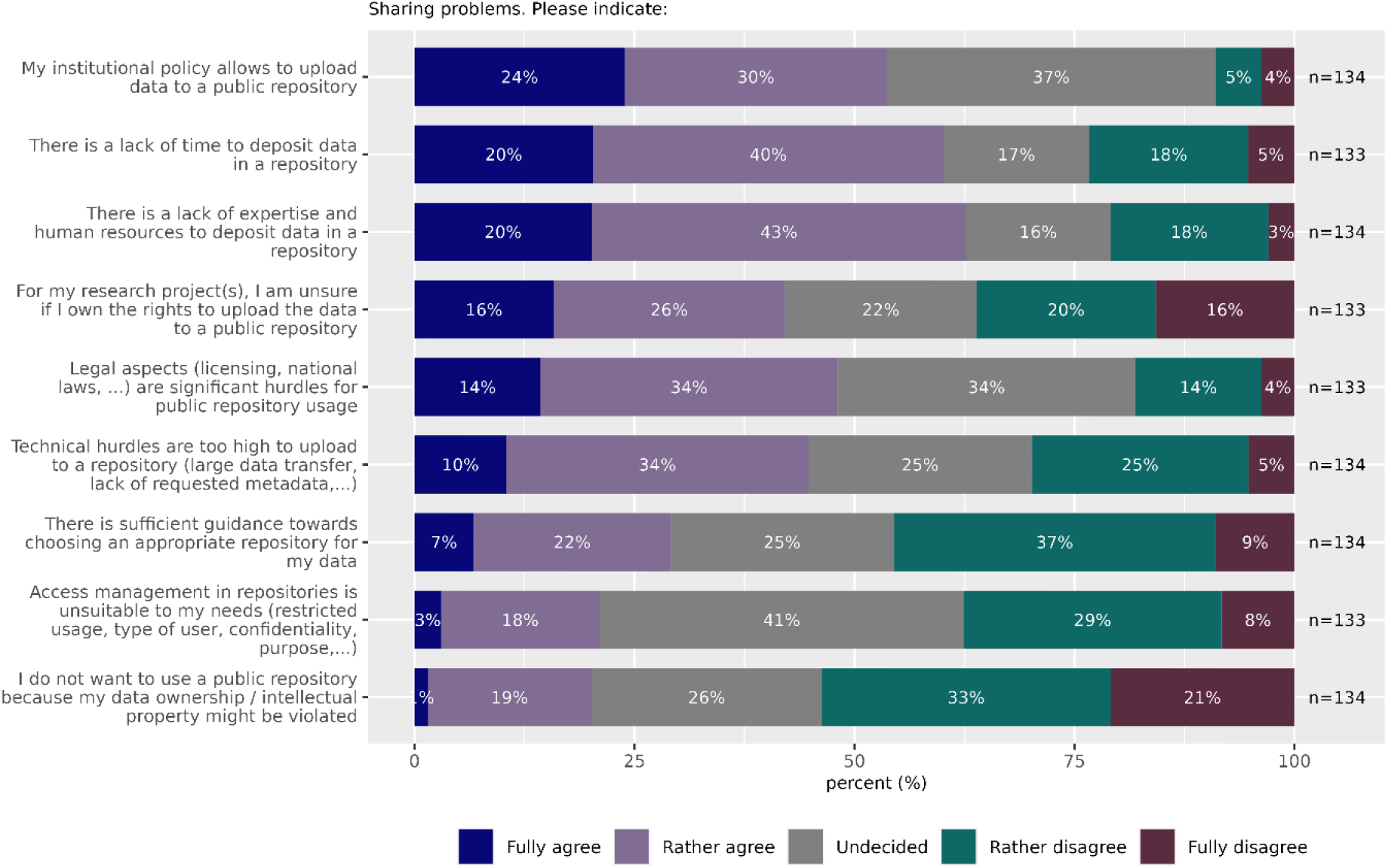
**Question 11** - Please choose the appropriate response for each item: Fully agree / Rather agree / Undecided / Rather disagree / Fully disagree

- My institutional policy allows to upload data to a public repository
- There is a lack of time to deposit data in a repository
- There is a lack of expertise and human resources to deposit data in a repository
- For my research project(s), I am unsure if I own the rights to upload the data to a public repository
- Legal aspects (licensing, national laws, …) are significant hurdles for public repository usage
- Technical hurdles are too high to upload to a repository (large data transfer, lack of requested metadata,…)
- There is sufficient guidance towards choosing an appropriate repository for my data
- Access management in repositories is unsuitable to my needs (restricted usage, type of user, confidentiality, purpose,…)
- I do not want to use a public repository because my data ownership / intellectual property might be violated

**Figure 13:**
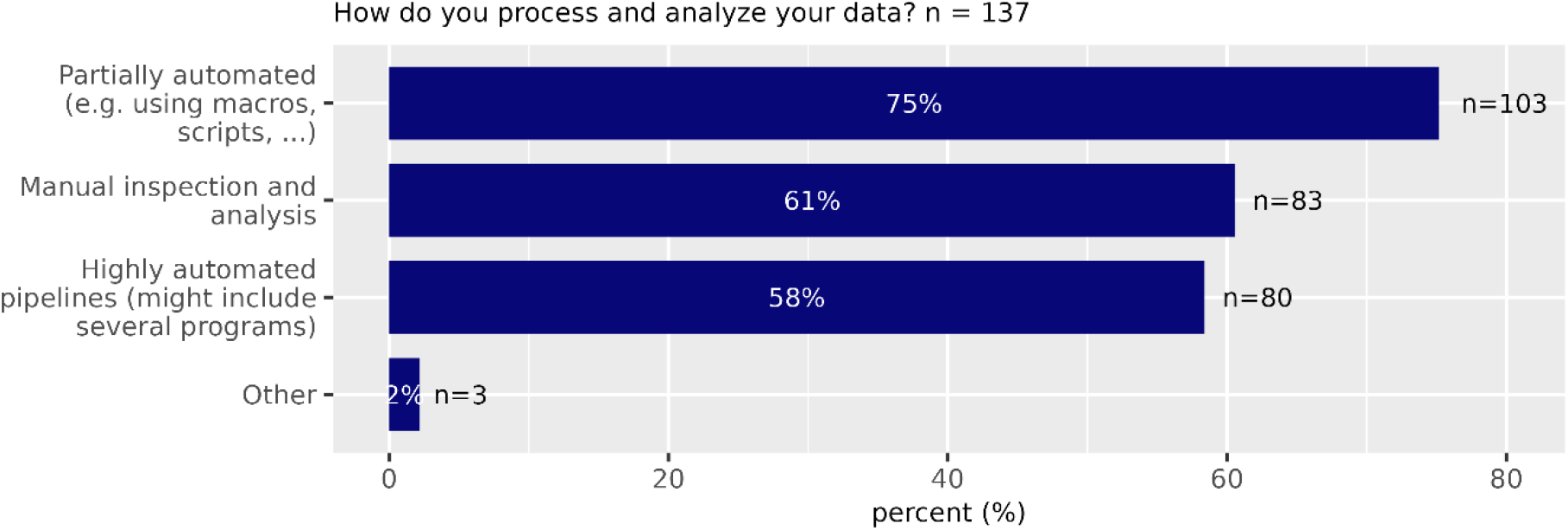
**Question 12** - How do you process and analyze your data? Please choose **all** that apply:

- Partially automated (e.g. using macros, scripts, …)
- Manual inspection and analysis
- Highly automated pipelines (might include several programs)
- Other:

**Figure 14:**
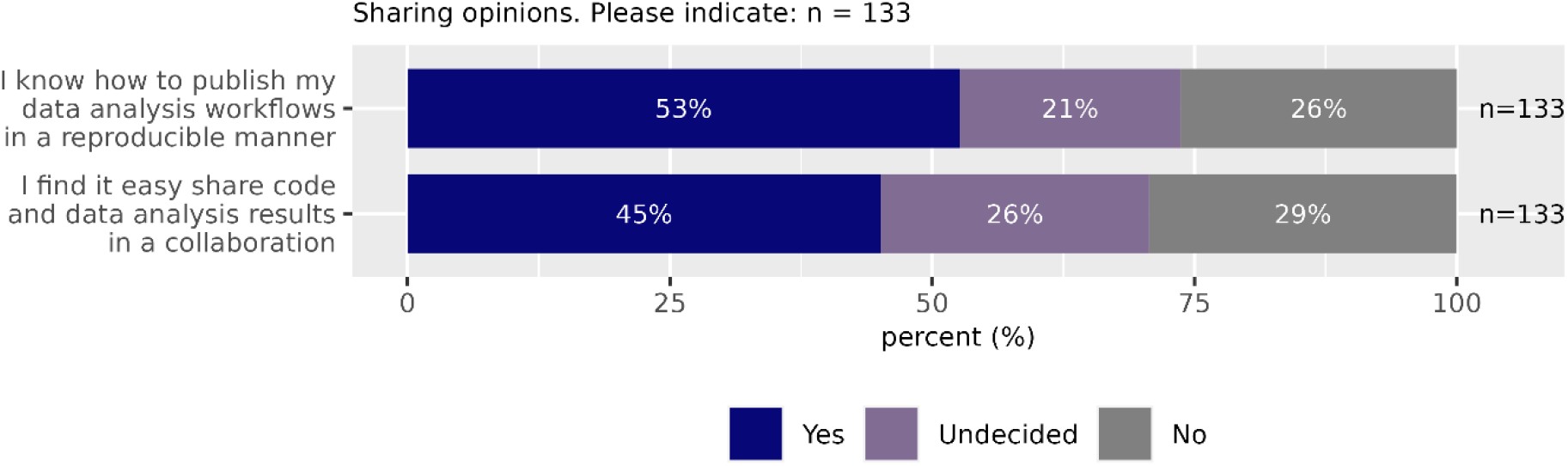
**Question 13** - Please choose the appropriate response for each item: Yes / No / Undecided

- I know how to publish my data analysis workflows in a reproducible manner
- I find it easy share code and data analysis results in a collaboration

### Own data management skills are largely seen as not being “high”

Research data management skills are essential for preparing, analysing, and publicly sharing data. 43% of responders disagree with the statement “Overall I am highly knowledgeable about research data management in my research field” (Fig. 15). Only 34% of the survey participants think that they have proficiency in research data management. Only 36% think they know which research data management methods are available, and 36% think they are “highly knowledgeable about research data management”. Interestingly, 59% of all respondents nevertheless agree or rather agree that they “can handle their research data according to community standards”. This could be due to the availability of data research managers who assist with data handling. However, only 19% of participants have dedicated personnel with research data management or data curation expertise in their labs.

**Figure 15:**
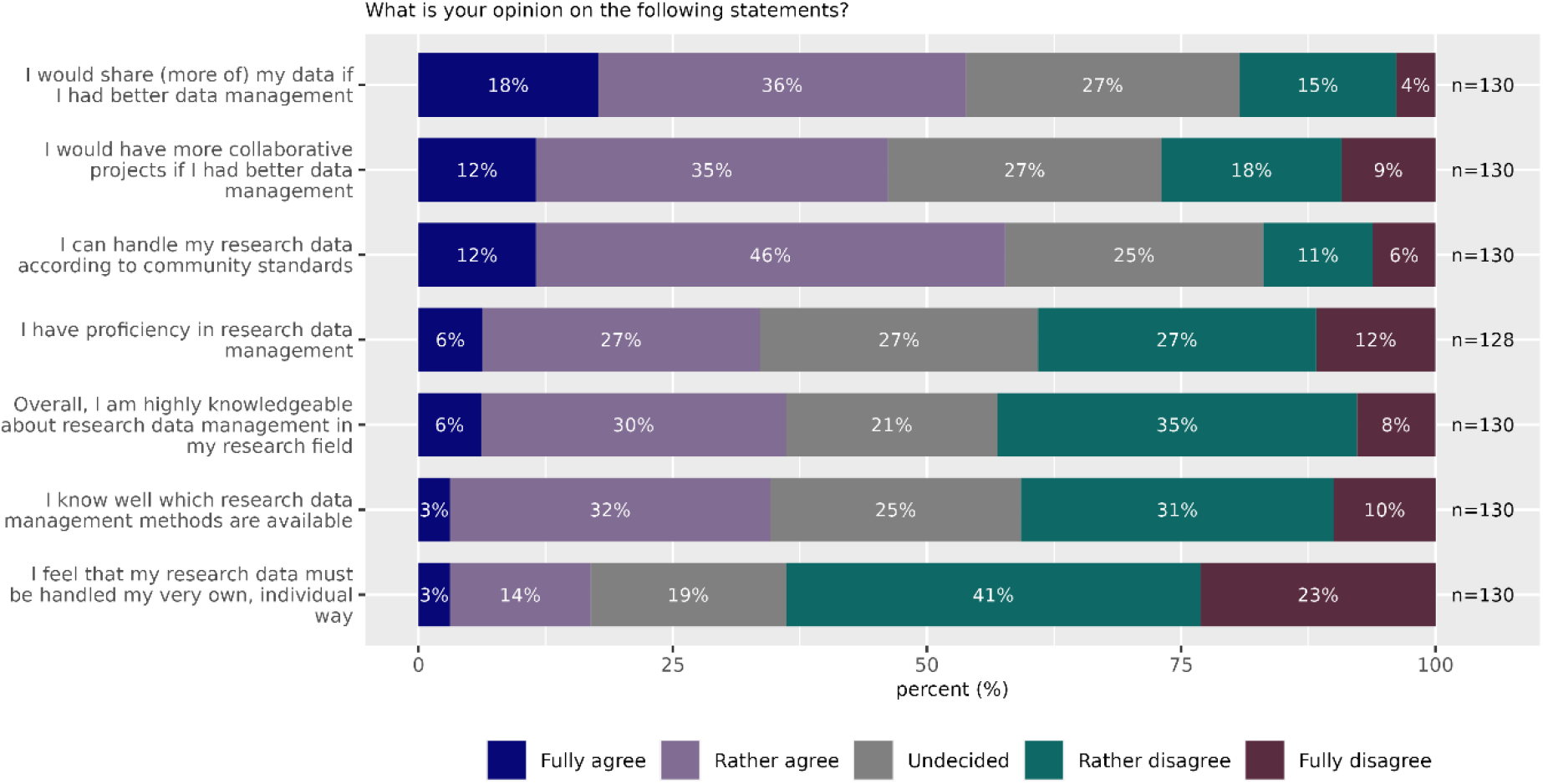
**Question 14** - What is your opinion on the following statements? Please choose the appropriate response for each item: Fully agree / Rather agree / Undecided / Rather disagree / Fully disagree

- I would share (more of) my data if I had better data management
- I would have more collaborative projects if I had better data management
- I can handle my research data according to community standards
- I have proficiency in research data management
- Overall, I am highly knowledgeable about research data management in my research field
- I know well which research data management methods are available
- I feel that my research data must be handled my very own, individual way

**Figure 16:**
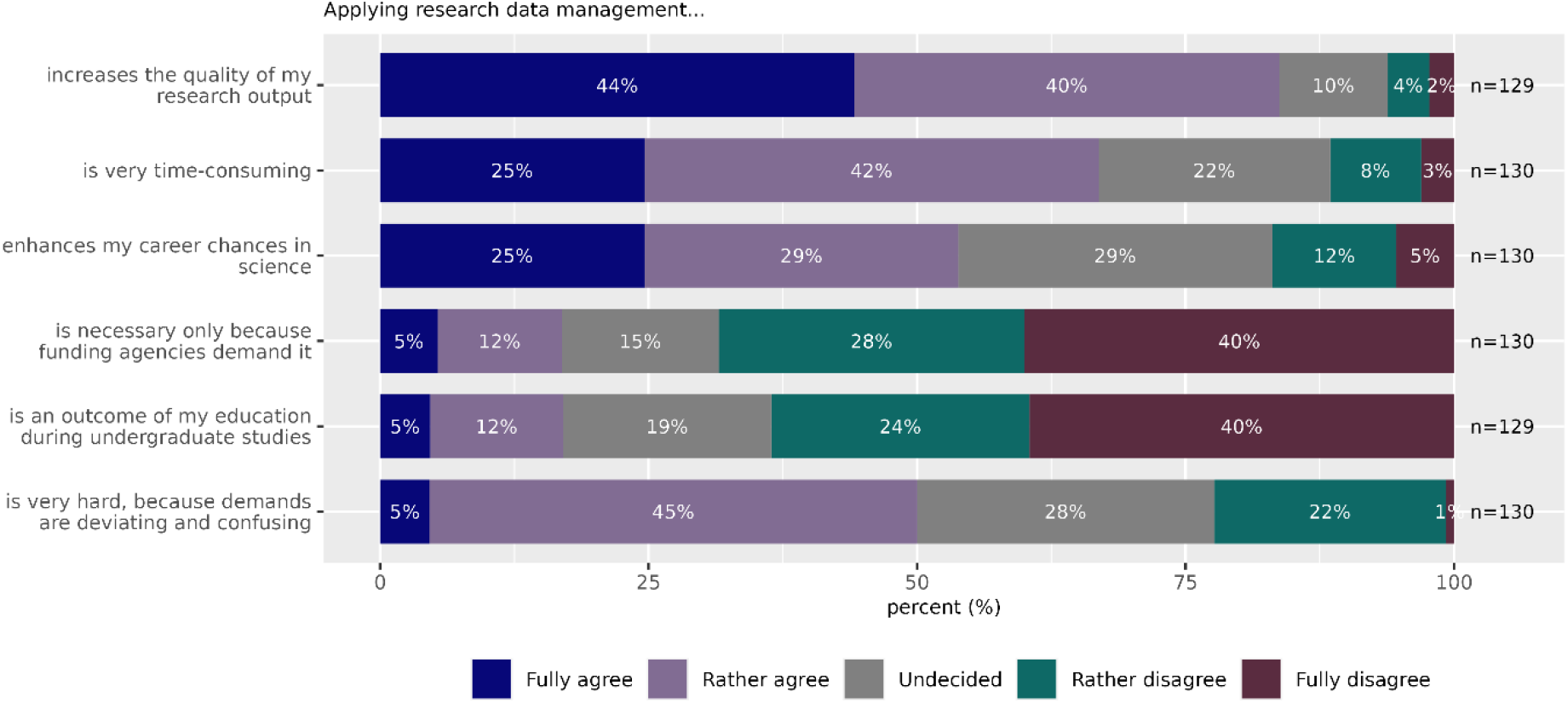
**Question 15** - Applying research data management…. Please choose the appropriate response for each item: Fully agree / Rather agree / Undecided / Rather disagree / Fully disagree

- increases the quality of my research output
- is very time-consuming
- enhances my career chances in science
- is necessary only because funding agencies demand it
- is an outcome of my education during undergraduate studies
- is very hard, because demands are deviating and confusing

**Figure 17:**
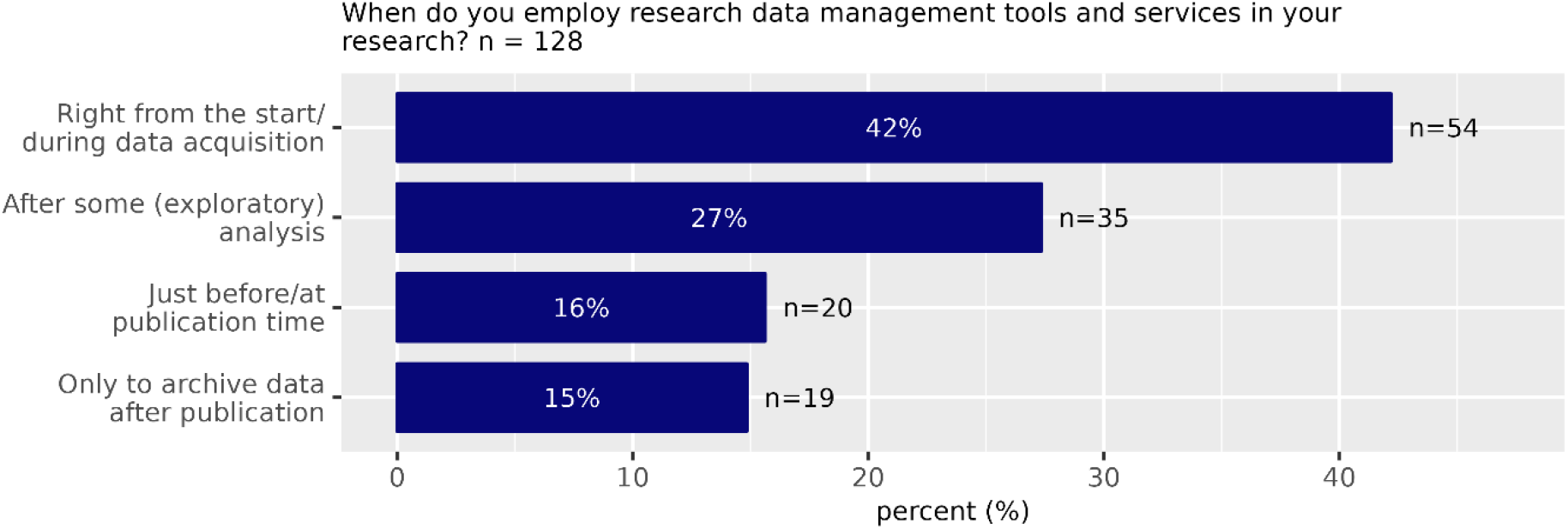
**Question 16** - When do you employ research data management tools and services in your research? Please choose **only one** of the following:

Right from the start/during data acquisition
After some (exploratory) analysis
Just before/at publication time
Only to archive data after publication

**Figure 18:**
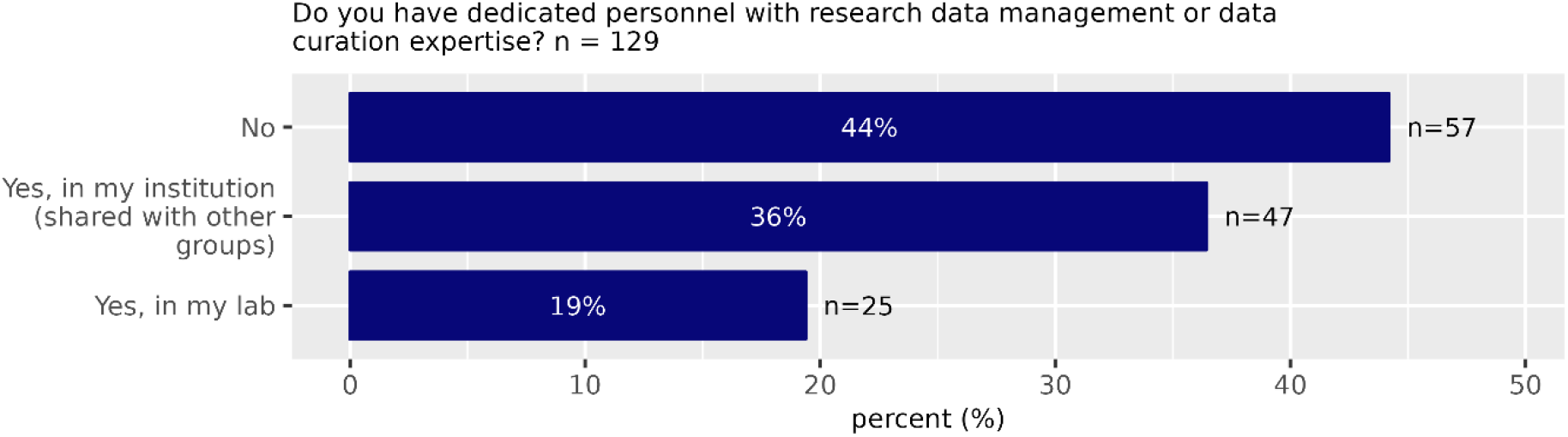
**Question 17** - Do you have dedicated personnel with research data management or data curation expertise? Please choose **only one** of the following:

No
Yes, in my institution (shared with other groups)
Yes, in my lab

**Figure 19:**
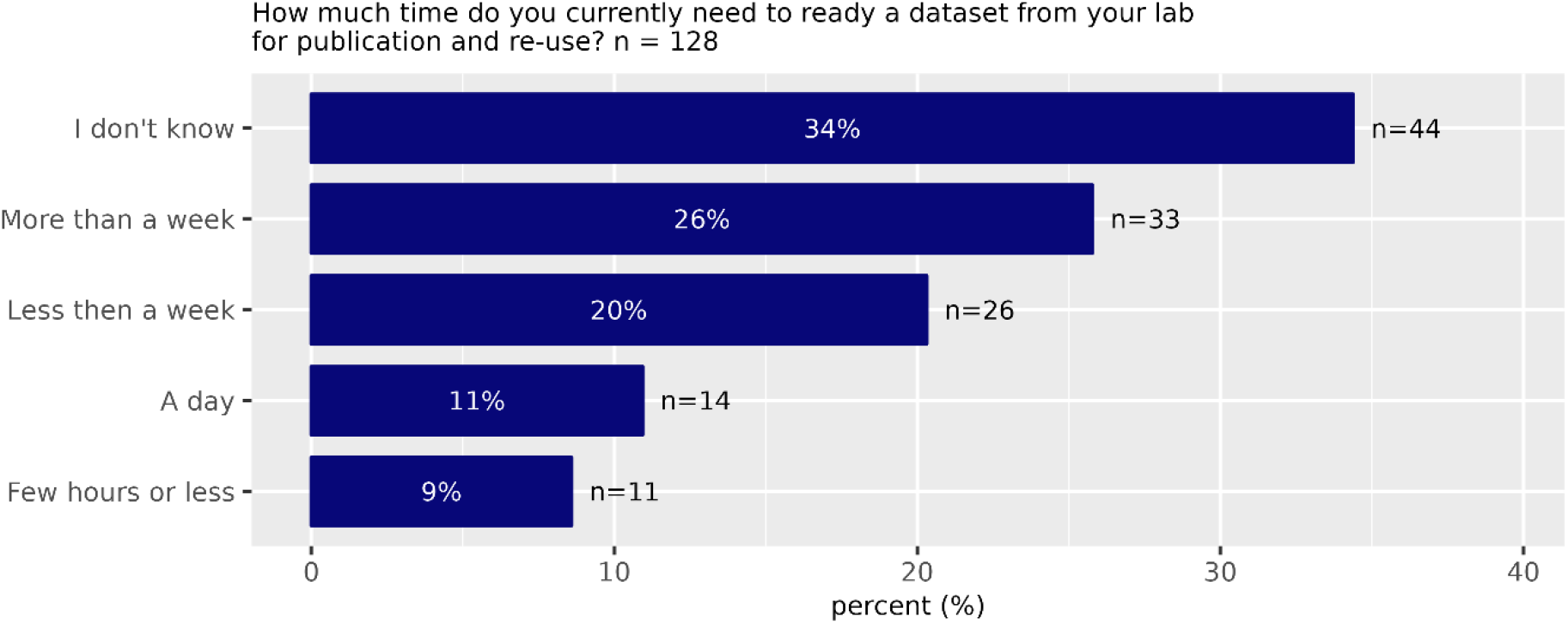
**Question 18** - How much time do you currently need to ready a dataset from your lab for publication and re-use? Please choose **only one** of the following:

- I don’t know
- More than a week
- Less then a week
- A day
- Few hours or less

We further investigated whether there is a dependency between public data sharing and the perception of one’s own competence regarding RDM. From the reported self-assessments, only the statement “I think that I can handle research data according to community standards” (Fig. 15) showed a strong connection to the likelihood that data are shared openly (Fig. 8). Participants agreeing to this statement were six times more likely to share data publicly than those who disagreeing. Self-assessed relatively high competence in the other RDM capabilities, leads to an increase in data sharing as well – although to smaller degrees (increase factor 1.2 (I know which research data management methods are available), increase factor 1.4 (“Overall, I am highly knowledgeable about research data management in my research field”) and increase factor 1.75 (“I have proficiency in RDM”)). Thus, in summary, better RDM knowledge leads to increased data sharing.

### Tools and standards for RDM are not yet widely used

While standard tools for data processing (data analysis) are widely used, the use of standard RDM tools for data sharing is significantly lower (sharing data openly, metadata collection and management, provenance tracking, Fig. 6).

### Those scientists who use more tools or standards are more likely to share their data

In the group that did not share their data publicly, only 33% use tools or standards, while in the group that share data, 54% use available tools or standards. A possible explanation could be that scientists who work a lot with standard tools find it easier to employ the rules and have a higher digital literacy required for the public sharing of data. Alternatively, the motivation to share data may be a strong driver to adopt standard methods. Respondents who share their data publicly are 42% more likely to use standard tools “mostly” in their daily work compared to respondents who did not share their data publicly.

### Perceived obstacles for research data management and sharing

Only 20% of the participants indicate a reluctancy to share data publicly because the data ownership or intellectual property might be violated. Interestingly, 37% of participants do not know whether their institutional policy allows uploading data to a public repository, while only 9% are confident that their institutions do not support this.

Further, 58% are not sure whether they own the rights to upload data from their own research project. 48% see legal aspects as significant hurdles for public repository usage. These answers indicate substantial uncertainties about legal issues. Indeed, only 18% think that legal aspects are no significant hurdles for public repository usage.

Only 29% of participants think there is sufficient guidance for choosing an appropriate repository for their data. 63% believe that there is a lack of expertise and human resources to deposit data in a repository. 45% think that the technical hurdles are too high to upload data to a repository.

83% of respondents do not think that their research data must be handled in an individual way that would not be easily compatible with existing standards, tools, or guidelines (Fig. 15). The lack of professional data management is reported as a problem. 70 (54%) participants think they would share more of their data if they had better RDM, while only 27% believe that better RDM would not increase the amount of their own data to share.

70% of those respondents who have previously prepared data for publication and re-use indicated that the time they need to ready a dataset requires more than a day, and 39% need even more than a week. Accordingly, 60% think there is a lack of time to deposit data in a repository. In comparison, only 23% do not believe that time is a problem for depositing data in a public repository (Fig. 12).

Questioned for the most pressing issues hindering research data management and public data sharing, there is a strong consensus (i.e., about 70% of respondents are rating these problems as one of the top three (Fig. 20)) for the following two statements

- “Inappropriately documented custom code in non-reproducible computational environments”
- “Poor standardization of metadata and derived data”

**Figure 20:**
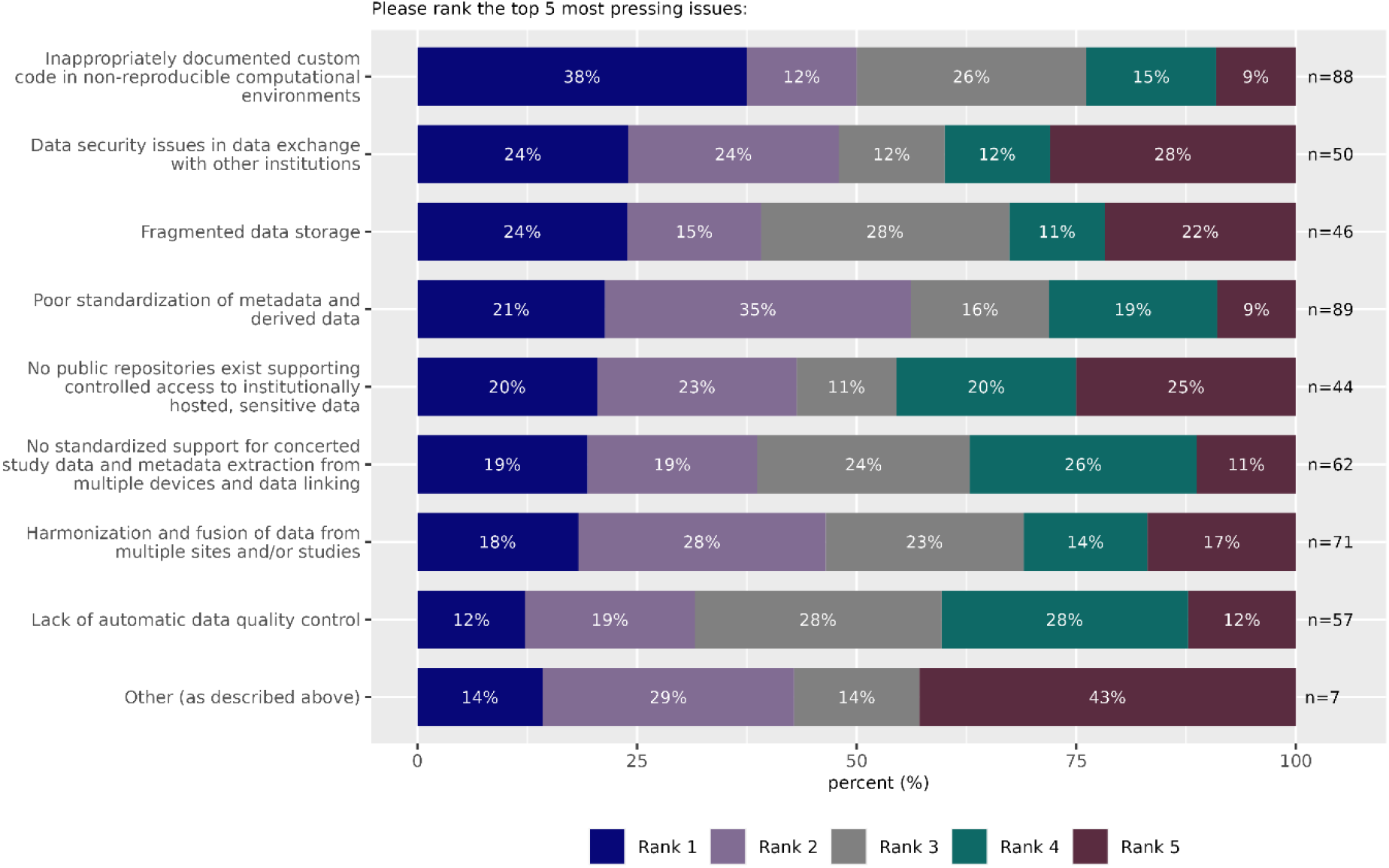
**Question 19** - Please rank the top 5 most pressing issues: All your answers must be different and you must rank in order. Please select at most 5 answers Please number each box in order of preference from 1 to 9 Please choose no more than 5 items.

- Inappropriately documented custom code in non-reproducible computational environments
- Data security issues in data exchange with other institutions
- Fragmented data storage
- Poor standardization of metadata and derived data
- No public repositories exist supporting controlled access to institutionally hosted, sensitive data
- No standardized support for concerted study data and metadata extraction from multiple devices and data linking
- Harmonization and fusion of data from multiple sites and/or studies
- Lack of automatic data quality control
- Other (as described above)

**Figure 21:**
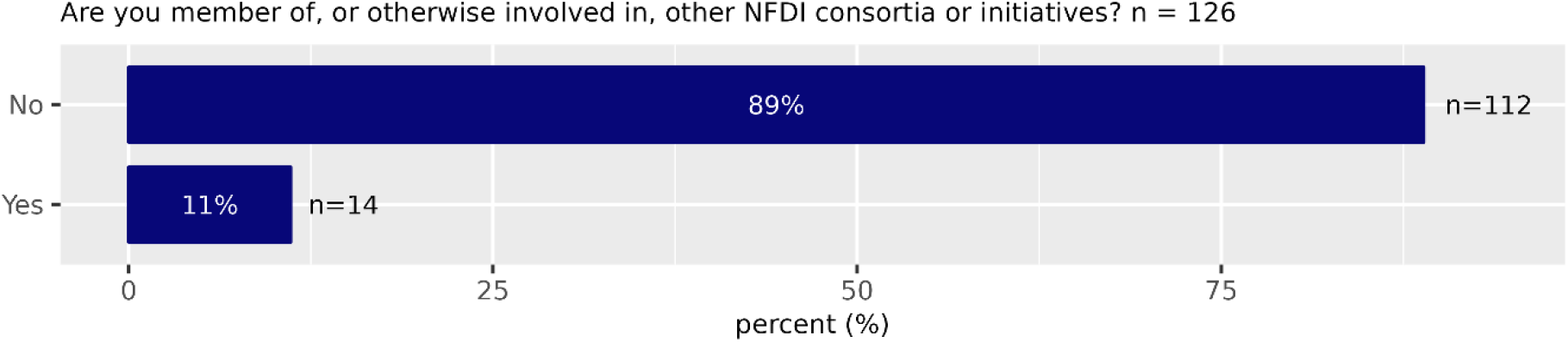
**Question 20** - Are you member of, or otherwise involved in, other NFDI consortia or initiatives? Please choose **only one** of the following

- No
- Yes

**Figure 22:**
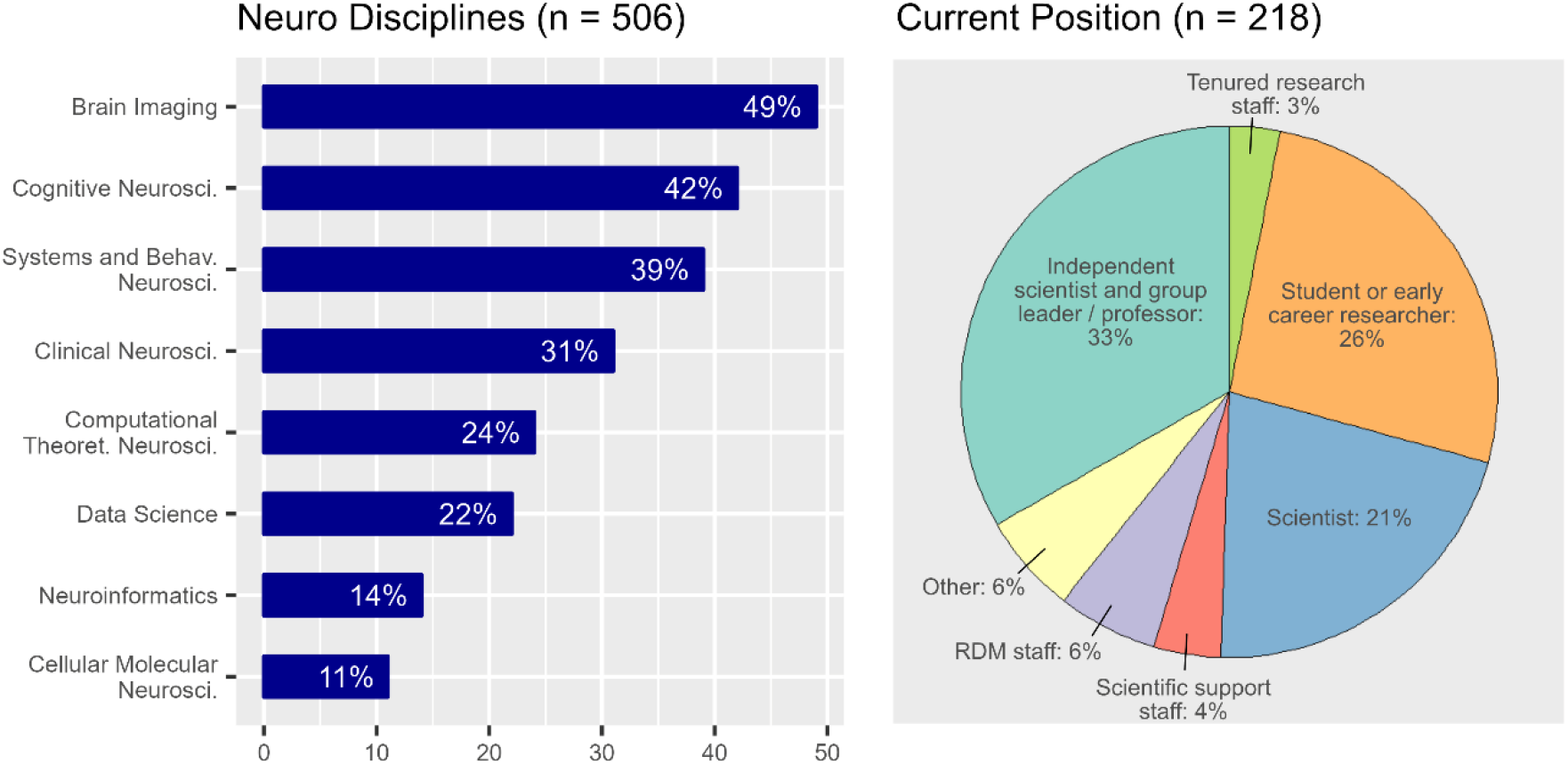
Distribution of neuroscience subdisciplines (multiple answers allowed, left), professional position of participants (right)

There are multiple concerns that are perceived similarly strong, but of lesser importance compared to the top two

- “Lack of automatic data quality control”
- “Harmonization and fusion of data from multiple sites and/or studies”
- “No standardized support for concerted study data and metadata extraction from multiple devices and data linking”
- “Data security issues in data exchange with other institution”

Also, general knowledge about methods and tools of research data management seems to be lacking. Only 34% of participants indicate that they think they know which RDM methods are available.

### Factors promoting public data sharing

To identify factors that promote public data sharing, we analysed separately the answers of the participants who had already shared their data in public repositories (n = 65). Thus, for this analysis we excluded all scientists who had never shared a single dataset publicly so far. The fraction of the different academic groups amongst scientists who have shared data publicly varied considerably: Whether data is shared depends on the position and experience of the person managing the data (Fig. 23).

**Figure 23:**
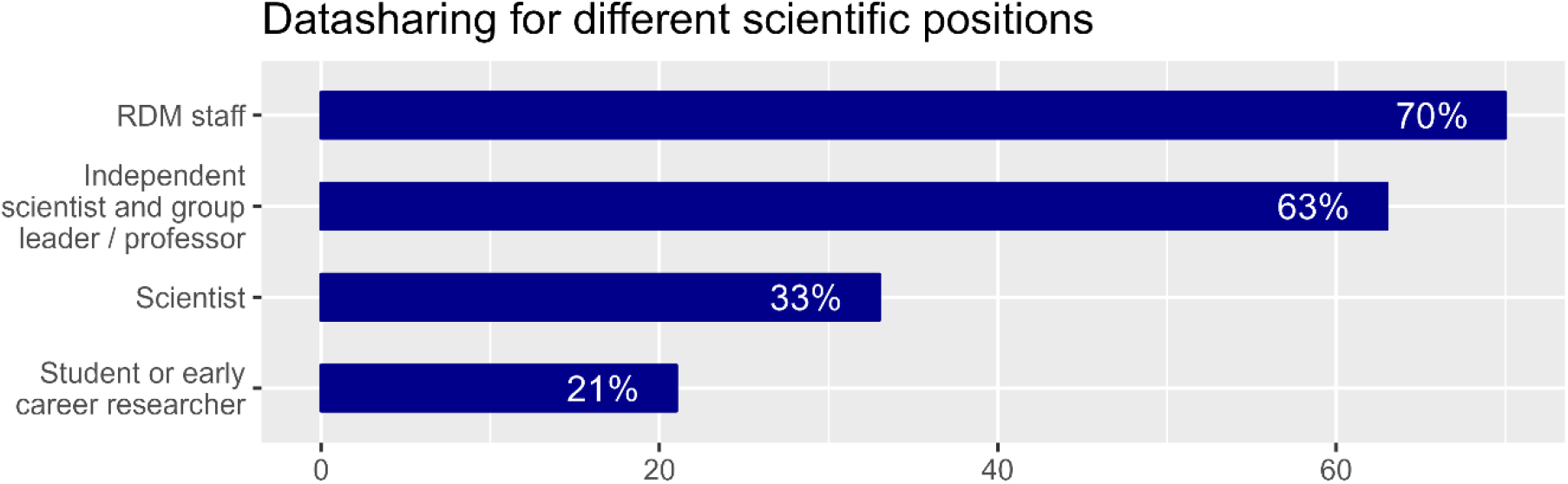
Percentage of respondents that have at least one dataset shared publicly shown separately according to their scientific position.

Interestingly, whether dedicated personnel with RDM or data curation expertise is available seems not to affect whether data are publicly shared. When dedicated RDM personnel is available, 56% of scientists report public data sharing, when it is not available (neither in the lab nor institution) 54% of scientists report public data sharing.

We analysed the dependence between the willingness to share data and the scientific sub-domain of the respective researcher. We found a relatively high degree of data sharing for scientists in the sub-domain of neuroinformatics, while we found a relatively low degree of data sharing in clinical neuroscience (Fig. 24).

**Figure 24:**
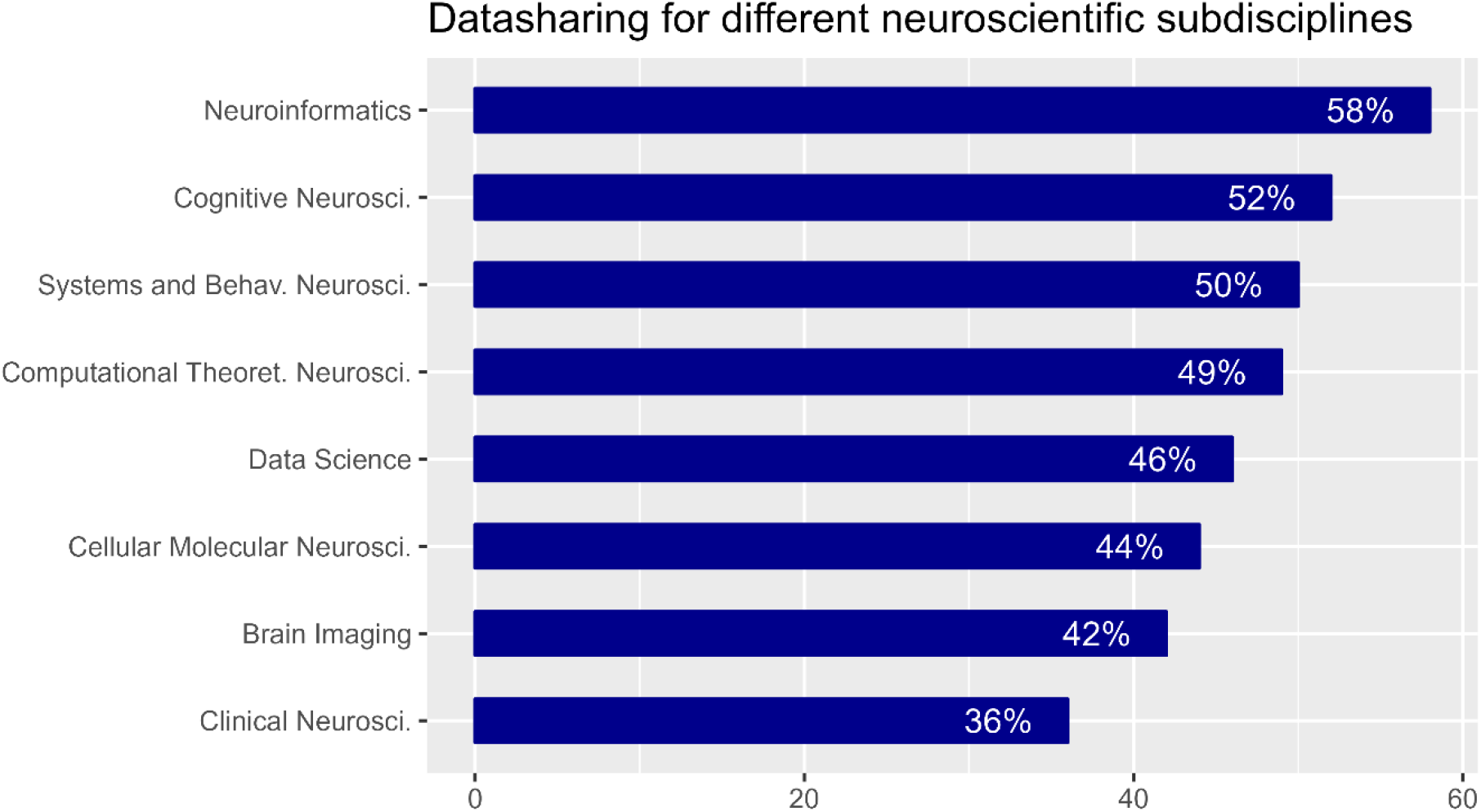
Percentage of respondents who have at least one dataset shared publicly - separated according to their scientific sub-domain.

## Discussion

### Work program of NFDI-Neuro based on the survey results

In their NFDI Cross-cutting Topics Workshop Report (Ebert, Barbara et al., 2021) the authors identified four prioritized areas where efforts in RDM are needed: (1) (Meta)Data, Findability, Terminologies, Provenance, (2) Research Data Commons, Infrastructure, Interoperability, Interfaces, Provenance, (3) Training & Education, (4) Ethical & Legal Aspects in General/Person related data management. Our survey confirms that these topics are perceived as of high relevance within the Neuroscience community.

The results of this survey led to the structure of NFDI-Neuro consortium work program with its five task areas

1. Community and Coordination – to provide training and coordination of activities
2. Data and metadata standards – to advance and disseminate existing standards
3. Provenance and workflows to advance and disseminate solutions for data lineage and digital reproducible workflows
4. Infrastructure and service – for data management and processing – including for sensitive data with Cloud and HPC resources
5. Modelling and big data analytics – for collecting RDM requirements from the perspective of the secondary data user community

NFDI-Neuro strives to address a major barrier hindering progress in neuroscience: the lack of integration of data and knowledge. The consortium plans to deliver a distributed and federated Virtual Research Environment (VRE) ecosystem for storing, sharing, analysing, and simulating complex neurobiological phenomena of brain function and dysfunction in a data protection compliant environment building on existing operational infrastructures for sensitive data at the Charité and EBRAINS. NFDI-Neuro will employ established technical and organizational principles to ensure lawful data management.

### NFDI-Neuro is built on achievements by the consortium that already serve thousands of researchers

The proposed NFDI-Neuro ecosystem builds on the experience of its leadership team and on the outcomes of successful previous infrastructure projects under their responsibility or with their participation such as the EBRAINS RI^8^ with its Health Data Cloud^9^ (Schirner et al., 2022), the European Open Science Cloud (EOSC) project Virtual Brain Cloud^10^ with its Virtual Research Environment^11^ (VRE) providing service for sensitive data to the research community.

### Data base development, data law and ethics

Coordinating institution Charité University Medicine Berlin (Charité) has served in the leadership of IT infrastructure projects for the last decade. Examples are: The Virtual Brian simulation platform^12^ with ca. 40k software downloads since its release and >150 publications of studies using the platform. The Virtual Research Environment, VRE^13^ - an operational and GDPR audited research infrastructure for sensitive data presently serving >90 users and >20 large-scale projects providing compute and storage resources via GUI and programmatic interfaces, data discoverability via a graph data base, access to high performance computing (HPC) and interfaces for data ingestion from hospital sources as well as support of the FAIR principles by accommodating existing data standards, lineage and provenance tracking. EOSC project Virtual Brain Cloud funded with €15Mill establishing infrastructure for sensitive data for the EOSC. The Health Data Cloud^14^ of EBRAINS RI serving more than 5000 registered EBRAINS users. As well as the just emerging eBRAIN-Health Horizon Europe Infrastructure project funded with €13M – a successor to EOSC Virtual Brain Cloud that establishes federated RI for sensitive health data. Charité holds recognized expertise in the field of data law and ethics underscored by their leading roles in large-scale projects developing GDPR compliant infrastructure for complex analysis of sensitive data and by Charité’s dedicated outreach activities on the topic e.g., the recent EU parliament workshop on digital data for dementia available as online resource^15^ or the GDPR Impact in Health Research Conference organized by EOSC Virtual Brain Cloud (led by Charité) with its contributions also being published online^16^ or publications in renowned journals such as on the topic “Brain Simulation as a Cloud Service” (Schirner et al. 2022) addressing technical and organizational measures implemented for the processing of sensitive health data in the cloud. Other partners hold similarly relevant activities and partnerships on an international scale - foremost EBRAINS AISBL - leading an EU RI funded with a billion Euros over the 10 years that recently got accepted to the ESFRI roadmap in a competitive selection process and thus will stay funded for the upcoming decades. Their work results in publications such as “The Human Brain Project: Responsible Brain Research for the Benefit of Society” (Salles et al. 2019).

### Generation of research data sets

NFDI-Neuro co-lead Research Center Julich (FZJ) contributes to DataLad^17^ and leads the DataLad data registry datasets.datalad.org that provides access to distributed datasets (0.5PB) across a wide range of repositories in a uniform fashion via DataLad. The FZJ Institute of Medicine alone has >1.2PB of data managed with DataLad in various databases.

NFDI-Neuro co-lead Fraunhofer Institute for Algorithms and Scientific Computing (SCAI) ADataViewer^18^ (Salimi et al., 2021) provides access to >20 large-scale cohorts with detailed information about the available data types and > 1200 mutually mapped data variables.

There are countless other examples how the NFDI-Neuro co-leads have engaged in the generation of research data sets – here we highlight a few: FZJ run https://www.studyforrest.org/ is operated for 8 years; Charité provides access to Hybrid Brain Model data^19^ (Schirner et al., 2018) available at Open Science Framework repository, various demonstration data sets for personalized brain simulation^20^ (Schirner et al., 2015) are also provided via the repository zenodo; time series and connectomes for personalized brain modeling in brain tumor patients^21^ (Aerts et al., 2018, 2020) are being made accessible via EBRAINS RI.

FZJ published an electrophysiological behavioral dataset of massively parallel recordings in macaque motor cortex (Brochier et al., 2018). Participant EBRAINS AISBL provides access to 978 curated data sets, 107 computational models, 166 software tools discoverable via knowledge graph.

### Data and metadata standards

NFDI-Neuro is involved in the leadership of data and metadata standards development. For instance Charité leads the BIDS (Brain Imaging Data Structure, (Gorgolewski et al., 2016) extension for Computational Modeling^22^ and has developed openMINDS metadata for computational models. FZJ has helped the development of the BIDS from its beginning as visible by the co-authorship on the seminal BIDS publication (Gorgolewski et al., 2016). It is also contributing to BIDS for electrophysiology (Holdgraf et al., 2019), the Neo data model (Garcia et al., 2014) – the latter also contributed to by NFDI-Neuro co-lead Ludwig Maximilian University (LMU) - and tools for the odML metadata standards (Sprenger et al., 2019). FZJ leads the development of OpenMINDS^23^ schema used by several hundreds of EU neuroscientists as indicated by more than 1793 contributors to the EBRAINS Knowledge Graph (KG)^24^ EBRAINS data collection – each item being made discoverable through openMINDS metadata annotation – be it a data set, software or a tool.

SCAI brings expertise to NFDI-Neuro in mapping scientific knowledge into computable, multi-scale mechanistic KGs (NeuroMMSig, Domingo-Fernández et al., 2017; Kodamullil et al., 2015), semantic frameworks, and in harmonizing datasets (ADataViewer). Further developments of SCAI include an Ontology lookup service^25^ and the Referential Ontology Hub for Applications within Neurosciences (Rohan)^26^.

### Complex interdisciplinary RDM work

The German neuroscience community is exemplary in its leadership in RDM and the development of provenance tools, semantic frameworks, metadata, and data standards. EBRAINS RI with scientific lead of FZJ and Sensitive Data Services lead Charite, the VRE, DataLad, openMINDS, SCAI ontologies and semantic frameworks, the German Neuroinformatics Node within the International Neuroinformatics Coordination Facility (INCF) with international training platforms and working groups for RDM. These achievements are under the lead of the NFDI-Neuro consortium and presently operational – supporting thousands of researchers in RDM. Already now, these solutions serve as blueprints for adoption by other research domains.

### NFDI-Neuro will establish a federated interoperable ecosystem for data and reproducible research

NFDI-Neuro’s proposed work includes the further advancement and dissemination of data and metadata standards, reproducible and interoperable digital workflows including container technology and DataLad, and infrastructure for FAIR (findable, accessible, interoperable and reusable; Wilkinson et al., 2016) sensitive data.

### Advancing previous RDM work

The proposed work comprises an ecosystem of data and metadata models with the capability for provenance tracking through version control and the executable storage of processes applied to the data with solutions like DataLad and container image technology and digital workflows for various types of data processing.

Building on the leading role of SCAI in the domain of semantic frameworks and ontologies we are convinced to achieve the deliverables as outlined. The challenges of integrating different semantic frameworks are being addressed successfully already and extensively in internationally leading roles by our partners, for instance by developing ontology lookup services and metadata mapping services. Efforts in the standardization of acquisition protocols and procedures are a crucial part of RDM – and of the proposed program.

### Making data FAIR requires the establishments of workflows for annotation, quality control, versioning

Data quality assurance has a broad set of diverse aspects of which selected ones that the consortium considers relevant are being addressed by the project. They include completeness of data sets, compliance to naming and data set structure standards, but also signal quality, presence of reference measurements, and accessory data acquisition for artefact correction procedures. Making the quality assurance workflows available as container images renders them reproducible and re-usable and in combination with DataLad the outcomes are fully versioned – a precondition for reproducible quality control.

There may be also workflows that require manual interaction. But even those can be made reproducible through combining the user decisions with DataLad as has been demonstrated in our recent publication on Brain Simulation as a Cloud Service (Schirner et al., 2022).

### Training and Dissemination

Training and dissemination activities by NFDI-Neuro include establishing working groups, transfer teams, workshops, interactive decision trees, demonstrators, and extensive educational material. While not yet funded, NFDI-Neuro has conducted in its preparation phase more than ten webinars that are available online, five community workshops with the presentations published online, and a Special Issue “NFDI – National Research Data Infrastructure” with four review articles (Denker et al., 2021b; Hanke et al., 2021; Klingner et al., 2021; Wachtler et al., 2021) and an editorial (Denker et al., 2021a).

In addition, NFDI-Neuro is conducting weekly or biweekly online community meetings^27^ for two years with an average attendance of >40 persons per event. The meeting series will continue and is open to interested community members and accessible via a self-registration option on the NFDI-Neuro website.

The collaboration with EBRAINS AISBL and INCF guaranties central discoverability of all training materials, e.g., via the INCF training space or EBRAINS portal. Another example of NFDI-Neuro activity serving and addressing the whole community are the planned dynamic support actions that provide flexible funds for the development of solutions for newly identified RDM use cases over the course of the project. It is also planned to establish and coordinate a network of CRC data managers and to develop an RDM curriculum to generate impact across the whole Neuroscience community.

### Use Cases

The NFDI-Neuro Use Cases for RDM development have been selected from nine existing large collaborative research centers (CRCs) across Germany – thus representing state-of-the-art requirements of the German neuroscience community. The Use Cases are cross cutting the various RDM domains covered in the work program of NFDI-Neuro.

### Ethics, data protection and sharing

NFDI-Neuro addresses ethical and legal challenges linked to the development and use of digital twins for research, including: a) Representation, bias, consent, inclusivity of research and promotion of diversity, b) legal capacity shared and supported decision-making, communication, trustworthiness, and trust. Related deliverables and milestones in the work program comprise e.g., trustworthy AI, and an AI Ethics Whitepaper.

Data protection measures comprise technical and organizational measures including the conclusion of various types of processing, sharing and service agreements for processing or storage of data in research infrastructures.

### NFDI

NFDI-Neuro is in active and continuous exchange with other NFDI consortia (Figure 25). NFDI-Neuro members also participate in the work of the four NFDI Sections^28^ – the focus domains of the NFDI Association^29^.

**Figure 25.**
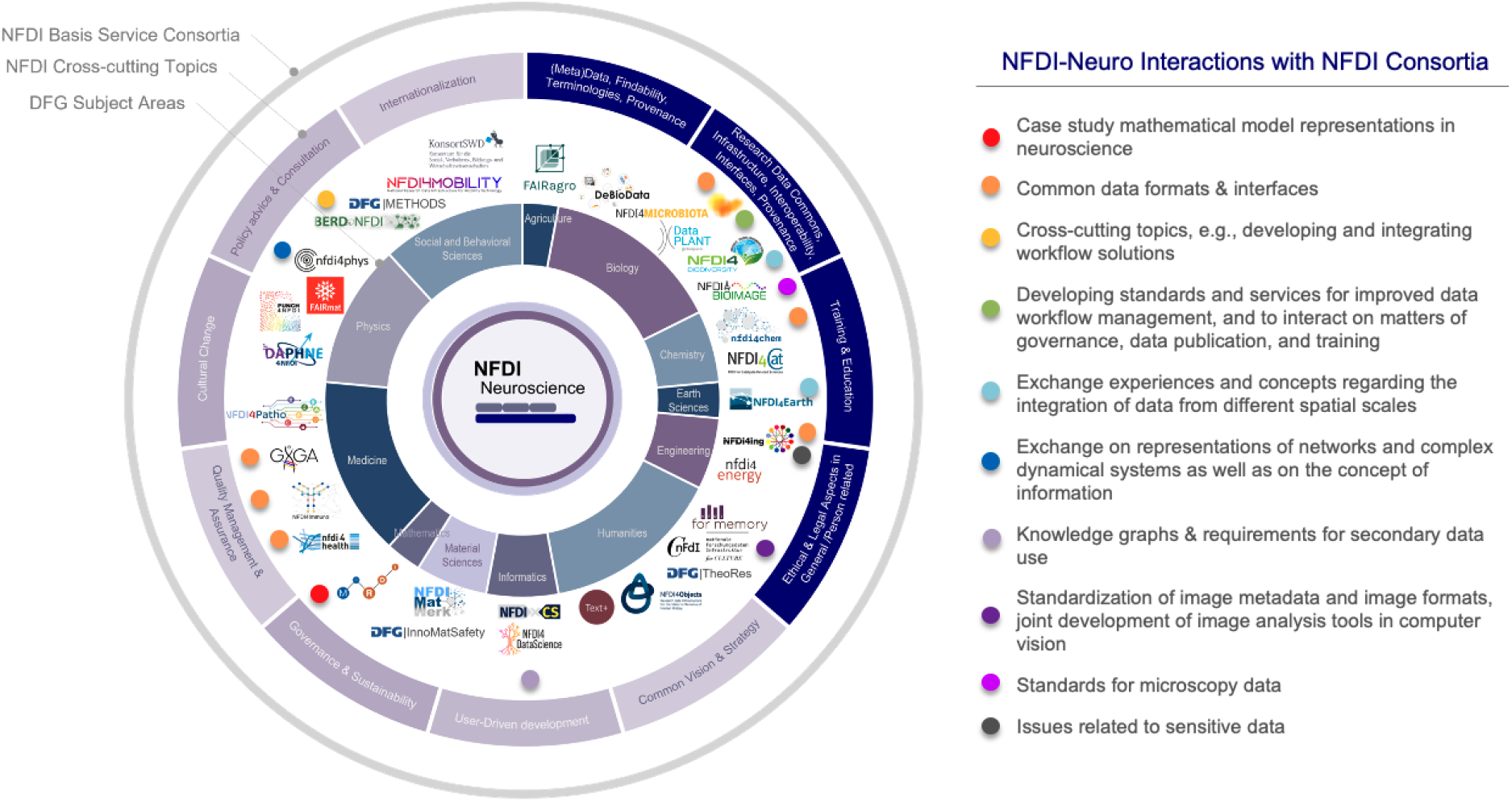
NFDI-Neuro has established several interactions with other NFDI consortia. Ten specific interactions points have bene identified as indicated on the right.

### International embedding and Sustainability

EBRAINS as an EU RI on the ESFRI roadmap will provide the means of making the developments of NFDI-Neuro sustainable and achieve international alignment and dissemination - thus ensuring international acceptance of all national developments. In addition, NFDI-Neuro members are leading ESOC and large-scale EU infrastructure projects for sensitive data services. They also participate in leadership of the international neuroinformatics initiative INCF that involves researchers of all continents and countries.

The tight collaboration with international long-term funded initiatives such as the INCF, EBRAINS on the ESFRI roadmap and the EOSC ensure sustainability of NFDI-Neuro activities.

### Value proposition for users outside NFDI-Neuro

NFDI-Neuro members take already now leading roles in RDM projects with existing large user communities benefiting from these activities. Dissemination includes activities such as the above-mentioned EU parliament Data sharing and GDPR workshops or workshops on digital twins with patient organizations and policy makers, or the international release of metadata schemas via INCF, Whitepaper Digital Twins, AI & Ethics, and various RDM Demonstrators.

### The role of the research community

Interestingly, in a recent Nature RDM survey (*Nature* **556**, 273-274 2018), 58% of participants think that the researchers hold the key role for improving the reproducibility of research and 91 % see them amongst the top three stakeholders to achieve this – thus being in a leading role ahead of laboratory heads, publishers, funders, department heads and professional societies who also were amongst the choices. This is in great alignment with what we experience in our work within RDM neuroscience projects (NFDI-Neuro, INCF, EBRAINS, central informatics projects of collaborative research centres funded by DFG (known as “INF projects”): It is the researchers themselves who are required to do the RDM and to co-develop RDM tools - and hence require training to obtain RDM literacy. “Reproducibility is like brushing your teeth. Once you learnt it, it becomes a habit” (Irakli Loladze in Baker et al. 2016). NFDI-Neuro aims to bring RDM to the individual labs – via several mechanisms including the establishment of transfer teams, working groups and massive training offers.

Together with the research community, NFDI-Neuro plans to co-design end-to-end services for storing, handling, annotating, and sharing complex neurobiological data, performing rigorous data processing and integration, and simulating neurophysiological phenomena. Data will be processed and annotated using common data models to ensure interoperability, re-usability and alignment with spatial and temporal reference frameworks. While an initial focus will be placed on selected Use Cases reflecting central projects of ongoing large-scale collaborative neuroscience projects across Germany, concepts of the NFDI-Neuro VRE ecosystem can subsequently be applied as a benchmark/model for other scientific foci in science and health areas. To this end, NFDI-Neuro aims the establishment of a community fostering the tight exchange between the science community and tool developers, supported by dedicated transfer teams that expedite the practical uptake of RDM tools in individual laboratories. The NFDI-Neuro infrastructure is designed to respect the sensitive nature of medical data – that includes for example brain scans or electroencephalograms (EEG), while making these data accessible for further research – with the proper technical and organizational measure in place to provide proper protection of subjects’ and patients’ rights. NFDI-Neuro will be tightly interlinked with major academic data producers in Germany, including the Collaborative Research Centers, multi-centre studies of the DZNE National Neuroimaging Network and the MII Initiative, thus enabling streamlined data deposition with full technical metadata transfer.

### Outlook

The survey builds the basis for a follow up world-wide survey presently developed by the International Neuroinformatics Coordination Facility (INCF) Infrastructure Committee.

## Conclusion

With the present survey, we identified various challenges in RDM in the neuroscience community. We found that the community perceives significant deficits with respect to transparent and reproducible data handling, annotation and sharing. Researchers with more experience and knowledge in RDM are more likely to share data for secondary use by their colleagues.

In summary:

- Only one third of neuroscientists think they have proficiency in RDM.
- Less than a quarter of the research teams have RDM staff.
- More than a third do not know if institutional policies allow loading data to a repository.
- Two third are not sure they own rights for uploading data to public repositories.
- Half of the researchers see legal hurdles for data sharing.
- Forty percent of those researchers who have previously prepared data for publication. and re-use say that the time they need to ready a dataset requires more than a week.
- Sixty percent think there is a lack of time to deposit data in a repository.
- Only one third think they know which RDM methods are available.

We are encouraged by the fact that only a minority of one fifth of respondents in the neuroscience community are not inclined to share data for secondary use and that literacy in the usage of tools and standards increases the frequency of data sharing. Thus, the survey results suggest that training, the provision of properly secure and protected research infrastructure, tools, standards, and additional resources for RDM are promising approaches to leverage RDM and foster reproducible and economic research in neuroscience. NFDI-Neuro will deliver on these topics. Therefore, we are convinced that we are addressing with NFDI Neuro the most pressing needs of our community. Our consortium has significantly contributed to several of the crosscutting goals of NFDI in the past. NFDI-Neuro plans to advance these operational solutions and to transfer them to an increasing number labs of the German and international science community.

Raw data of the survey has been published in the research data repository https://gin.g-node.org/NFDI-Neuro/SurveyData under the DOI 10.12751/g-node.w5h68v.

## Acknowledgment

PR gratefully acknowledges support by H2020 Research and Innovation Action grants Human Brain Project SGA2 785907, SGA3 945539, VirtualBrainCloud 826421, European Innovation Council grant PHRASE 101058240 and ERC 683049; Berlin Institute of Health & Foundation Charité, Johanna Quandt Excellence Initiative, German Research Foundation SFB 1436 (project ID 425899996); SFB 1315 (project ID 327654276); SFB 936 (project ID 178316478); SFB-TRR 295 (project ID 424778381); SPP Computational Connectomics RI 2073/6-1, RI 2073/10-2, RI 2073/9-1. MH was supported by the German Federal Ministry of Education and Research (BMBF 01GQ1905), Human Brain Project SGA3 945539, German Research Foundation SFB 1451 (project ID 431549029); GRK 2150 (project 269953372). MD, SG are supported by Human Brain Project SGA3 945539 and the Helmholtz Metadata Collaboration (HMC). SOJ is supported by the Federal State of Saxony-Anhalt, Germany (FKZ: I 88).

## Footnotes

5 https://vre.charite.de/xwiki/wiki/vrepublic/view/Main/VRE_Community/community_meetings/

6 https://nfdi4bioimage.de/

7 https://gin.g-node.org/NFDI-Neuro/SurveyData (https://doi.org/10.12751/g-node.w5h68v)

8 Ebrains.eu

9 https://www.healthdatacloud.eu/

10 https://cordis.europa.eu/project/id/826421

11 https://vre.charite.de/vre

12 thevirtualbrain.org

13 https://vre.charite.de

14 https://www.healthdatacloud.eu/

15 https://www.youtube.com/playlist?list=PLO-PgQHI1WQX6SZ44tR2SCi9b9xOVrv37

16 https://www.youtube.com/watch?v=LcTCF8veshc

17 https://www.datalad.org/

18 https://adata.scai.fraunhofer.de/

19 http://dx.doi.org/10.17605/OSF.IO/MNDT8

20 https://zenodo.org/record/3497545#.YjcIo5rMJTZ

21 DOI: 10.25493/1ECN-6SM

22 https://bids.neuroimaging.io/get_involved.html#extending-the-bids-specification

23 https://github.com/HumanBrainProject/openMINDS

24 https://search.kg.ebrains.eu/

25 https://ols.neuro.scaiview.com/ontologies

26 https://rohan.scai.fraunhofer.de

27 https://vre.charite.de/xwiki/wiki/vrepublic/view/Main/VRE_Community/community_meetings

28 https://www.nfdi.de/sections/?lang=en

29 https://www.nfdi.de/?lang=en

